# Substrate specificity in lipoylation pathways: How lipoate:protein ligases discriminate between protein substrates

**DOI:** 10.64898/2026.10.01.755906

**Authors:** Carolin Kümpel, Emma Holz, Tomohisa Sebastian Tanabe, Marc Gregor Mohr, Christiane Dahl

## Abstract

Lipoate is an essential protein-bound cofactor attached to conserved lysine residues in structurally related acceptor proteins, including GcvH of the glycine cleavage system and LbpA proteins required for sulfur oxidation by the sulfur-oxidizing heterodisulfide-like (sHdr) complex. The coexistence of distinct lipoylation pathways and acceptor proteins raises the question of how the enzymes initiating lipoate attachment discriminate between homologous but functionally distinct substrates. We systematically tested lipoate:protein ligases and potential acceptor proteins from five representatives of the bacterial phyla Pseudomonadota and Aquificota using native gel-shift assays and MALDI-TOF mass spectrometry. The canonical *Escherichia coli* lipoate:protein ligase Lpl(AB) preferentially modified GcvH proteins but showed little or no activity toward LbpAs. In contrast, sulfur oxidation-associated lipoate:protein ligases, termed sLpl(AB), which function in the sLpl(AB)–LipS1/LipS2 lipoate assembly pathway, strongly preferred LbpA proteins and generally failed to modify GcvH. Conserved cysteine residues characteristic of LbpAs were dispensable for recognition by sLpl(AB). Instead, comparison of amino acid sequences and electrostatic surfaces revealed predominantly acidic interaction surfaces in GcvH proteins and corresponding basic regions in LbpAs. Accordingly, replacement of selected basic regions in *Roseovarius mucosus* LbpA2 by acidic residues reduced modification by its cognate sLpl(AB) and simultaneously enabled weak modification by *E. coli* Lpl(AB). These findings establish electrostatic surface properties as major determinants of substrate discrimination by lipoate ligases and show that recognition depends on distributed features of the folded acceptor protein rather than solely on the sequence surrounding the lipoylated lysine.

**SIGNIFICANCE STATEMENT:** Post-translational modification requires enzymes to distinguish their physiological targets from structurally similar proteins, yet the molecular principles underlying such discrimination often remain unclear. Here, we show that two evolutionarily distinct families of lipoate:protein ligases discriminate between the homologous acceptor proteins GcvH and LbpA, with major functions in central metabolism and sulfur oxidation, respectively. Substrate specificity is not determined by conserved cysteine residues or solely by the sequence surrounding the modified lysine. Instead, electrostatic properties distributed across the protein surface emerge as major determinants of recognition. Altering these properties shifts ligase preference, demonstrating how related protein substrates can be selectively directed toward parallel modification pathways.

## 1. INTRODUCTION

The cofactor lipoate (α-lipoic acid; 1,2-dithiolane-3-pentanoic acid, or thioctic acid) is an eight-carbon organosulfur compound that is essential across all domains of life and plays central roles in energy-conserving pathways, as exemplified by the lipoate-dependent pyruvate and α-ketoglutarate dehydrogenase complexes or the sulfur-oxidizing sHdr-LbpA system (Cao, Koch et al., 2018; Cronan, 2016; Cronan, Zhao, & Jiang, 2005; Rowland, Snowden, & Cristea, 2018; Spalding & Prigge, 2010). Lipoic acid is covalently attached to its target proteins through an amide bond linking its carboxyl group to the ε-amino group of a conserved lysine residue. Its characteristic 1,2-dithiolane ring contains sulfur atoms at positions C6 and C8, which confer redox activity and contribute to the antioxidant properties of lipoate (Acosta, Albers et al., 2025; Cussiol, Alegria, Szweda, & Netto, 2010; Nyvltova, Smutna, Tachezy, & Hrdy, 2016; Perham, 2000; Spalding & Prigge, 2010). Beyond its redox activity, the lipoyl prosthetic group is distinguished by its relatively large size and high flexibility compared with modifications such as phosphorylation or acetylation, allowing it to function as a mobile “swinging arm.” These flexible arms can span distances of up to approximately 3 nm, allowing the lipoyl group to shuttle covalently bound catalytic intermediates between successive active sites and, in some cases, between different subunits of multienzyme complexes (Perham, 2000). The functional role of lipoamide is particularly well exemplified by GcvH proteins, which typically comprise ~130 amino acids, have a molecular mass of ~15 kDa, and essentially consist of a single lipoyl domain forming the central component of the glycine cleavage complex (Kikuchi, Motokawa, Yoshida, & Hiraga, 2008; Okamura-Ikeda, Hosaka et al., 2010; Spalding & Prigge, 2010). The GcvH lipoyl moiety undergoes a cycle of reductive methylamination catalyzed by GcvP, methylamine transfer catalyzed by GcvT, and reoxidation catalyzed by GcvL (dihydrolipoamide dehydrogenase, DLDH) (Spalding & Prigge, 2010). Thus, lipoate serves as an electrophile that transiently binds reaction intermediates, as a flexible swinging arm that channels substrate between the active sites of different enzymes, and as a redox-active cofactor that cycles between reduced and oxidized states.

In 2-oxo acid dehydrogenases, the lipoyl group is attached to a lysine residue in one or more independently folded lipoyl domains of ~80 aa amino acid residues (Jones, Horne, Reche, & Perham, 2000; Jones, Stott, Howard, & Perham, 2000; Perham, 2000). The lipoyl domain consists of an eight-stranded β-barrel consisting of two four-stranded anti-parallel β-sheets. The lipoyl–lysine residue is prominently displayed in a tight β-turn. The overall structure of GcvH is highly similar, with the lipoyl-lysine residue prominently exposed in a surface-accessible β-turn (Brocklehurst & Perham, 1993; Fujiwara, Maita et al., 2010; Pares, Cohen-Addad et al., 1994) (Fig. 1). The lipoate-binding LbpA proteins are closely related to GcvH proteins but form a distinct phylogenetic group that can be divided into two major clades, LbpA1 and LbpA2 (Cao, Koch, et al., 2018). Despite their phylogenetic divergence, LbpA proteins share the characteristic fold of GcvH and the same conserved site for covalent lipoate attachment (Fig. 1). They are found in diverse dissimilatory sulfur-oxidizing bacteria that use reduced sulfur compounds as electron donors for energy conservation, where they are essential for the cytoplasmic oxidation of sulfane sulfur to sulfite via the sulfur-oxidizing heterodisulfide reductase-like (sHdr) complex (Cao, Koch, et al., 2018; Koch & Dahl, 2018). Up to three distinct LbpA proteins can be encoded within *shdr-lbpA* gene clusters (Kümpel, Grosser, Tanabe, & Dahl, 2024; Mohr, Tanabe, Grosser, & Dahl, 2026; Tanabe, Grosser et al., 2023) (Fig. 2).

**FIGURE 1.**
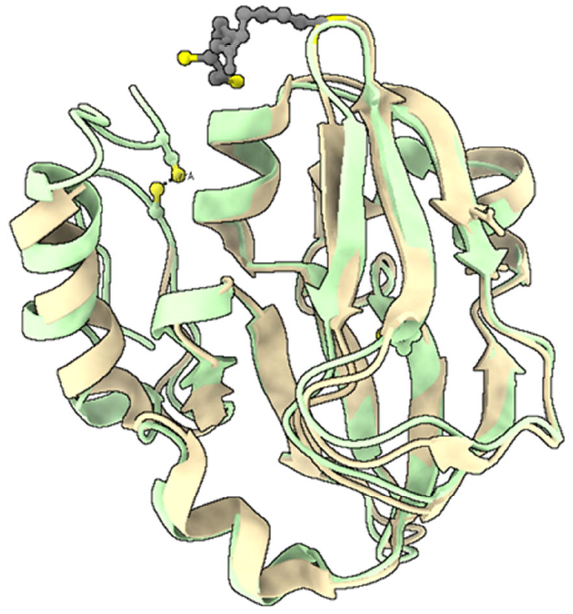
Superposition of *E. coli GcvH and LbpA2 from Hyphomicrobium denitrificans*. The structure of lipoylated *E. coli* GcvH (PDB 3AB9) is shown in beige and was determined by X-ray crystallography at 2 Å resolution (Okamura-Ikeda et al., 2010). The LbpA2 protein from *H. denitrificans* (Hden_0696) is essential for thiosulfate oxidation (Cao, Koch, et al., 2018), was modelled by AlphaFold (Abramson, Adler et al., 2024) and is depicted in green. Lipoamide is shown in grey with its two sulfur atoms highlighted in yellow. The sulfur atoms of two cysteine residues close to the N- and C-termini of the hyphomicrobial protein are also shown. The distance between the sulfur atoms is 1.77 Å, close enough to permit formation of an intramolecular disulfide bond in the oxidized state (Sun, Wang et al., 2017).

**FIGURE 2.**
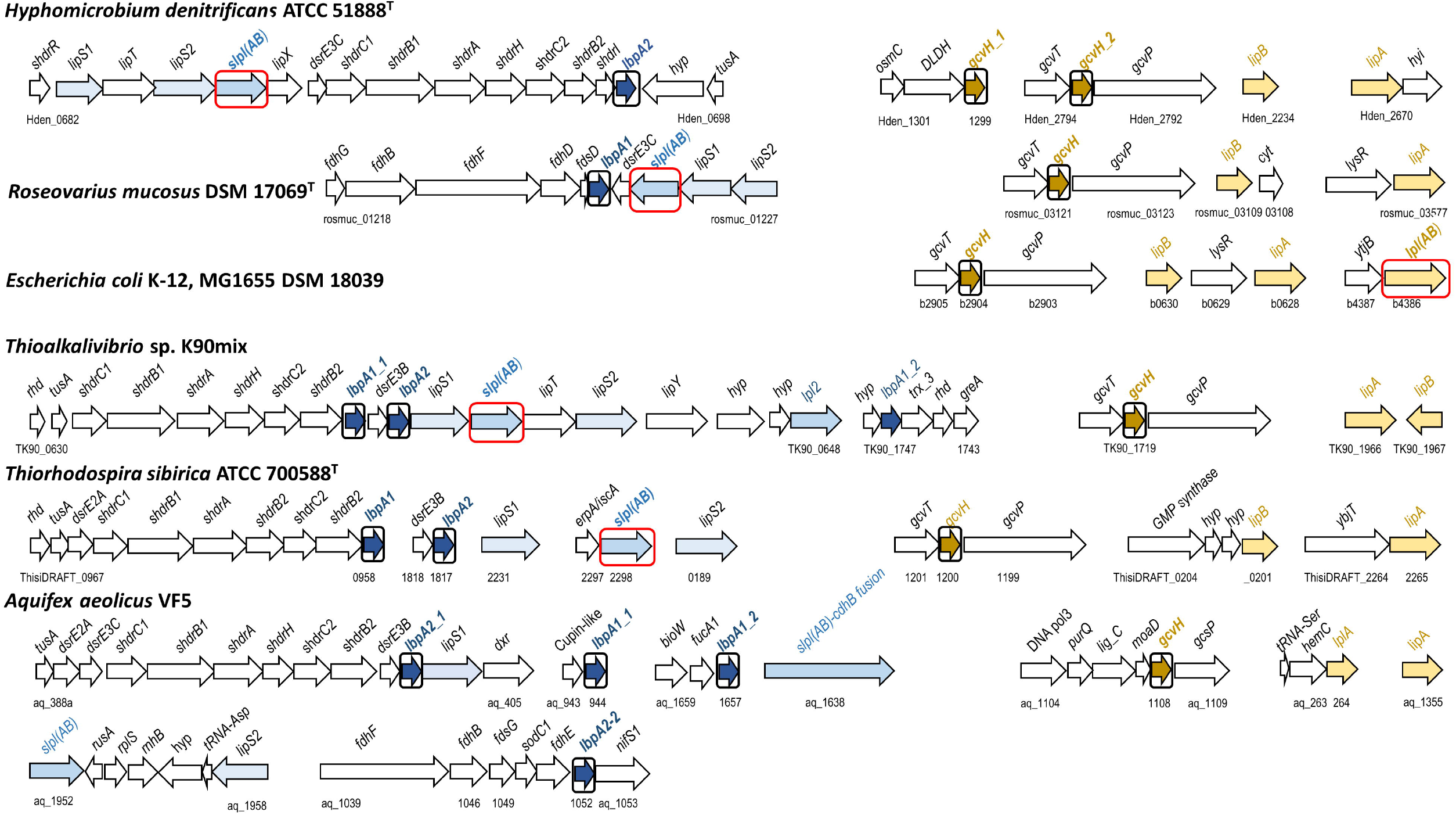
Arrangement of genes for lipoate assembly pathways and lipoylation substrates in five members of the Pseudomonadota and one representative of the Aquificota. The genes for 18 lipoylation substrates and the 5 lipoate:protein ligases studied in this work are highlighted by black and red boxes, respectively. For *Ts. sibirica* locus tags are given according to JGI-IMG. The *shdr* genes encode the cytoplasmic sulfur-oxidizing sHdr complex and accessory proteins. LipT is an FAD-binding oxidoreductase possibly aiding the LipS1/S2-catalyzed sulfur insertion step. *lipY*, putative long chain fatty acid transporter; *lipX*, putative glutamine amidotransferase. TusA, Rhd, DsrE, and MoaD proteins act as sulfur carriers/transferases. sLpl(AB), Lpl(AB), sLplA: lipoate:protein ligases. LipA, lipoyl synthase; LipB, lipoyl(octanoyl)transferase; LbpA, lipoate-binding protein; GcvH, glycine cleavage system protein H; DLDH, dihydrolipoamide dehydrogenase. The following annotations were adopted from JGI/IMG: *bioW*, 6-carboxyhexanoate-CoA ligase involved in biotin biosynthesis; *cdhB*, CO dehydrogenase beta subunit/acetyl-CoA synthase epsilon subunit; *cyt*, cytochrome *c*; DNA pol3, DNA polymerase III subunit delta; *dxr*, 1-deoxy-D-xylulose-5-phosphate reductoisomerase; *erpA/iscA*, iron-sulfur cluster assembly accessory protein; *fdh* and *fds*, subunits of formate dehydrogenase and accessory proteins; *fucA1*, L-fucolose-phosphate aldolase; *gcvT*, glycine cleavage system protein T (aminomethyltransferase); *gcvP* and *gcsP*, glycine cleavage system protein P (glycine dehydrogenase); *greA*, regulator GreA/B family; *hemC*, hydroxymethylbilane synthase involved in heme biosynthesis; *hyi*, hydroxypyruvate isomerase; *hyp*, hypothetical; *lig_C*, C-terminal region of RNA ligase, *lys*R, LysR-type regulator; *nifS1*, cysteine desulfurase; *osmC*, protein potentially involved in combatting reactive oxygen species (Nyvltova et al., 2016); *purQ*, glutamine amidotransferase domain; *rnhB*, ribonuclease HII; *rplS*, large subunit ribosomal protein L19; *rusA*, endodeoxyribonuclease; *sodC1*, superoxide dismutase; *trx*, thioredoxin; *ybjT*, uncharacterized protein.

Lipoylation is a rare but crucial post-translational modification, with both the enzymes that catalyze it and many of its target proteins evolutionarily conserved from bacteria to humans (Cronan et al., 2005; Lin, Chin et al., 2024; Rowland et al., 2018). Classically, two strategies for protein lipoylation have been distinguished: salvage of exogenous lipoate or its precursors and de novo synthesis from intermediates of fatty acid biosynthesis (Cronan, 2016, 2024). In bacteria, de novo lipoate synthesis is more common than lipoate scavenging (Tanabe et al., 2023). In the canonical salvage pathway, using free precursors involves lipoate:protein ligase Lpl(AB) that activates the precursors to lipoyl-or octanoyl-AMP at the expense of ATP before transfer to the target protein. In the canonical de novo pathway, an octanoyl transferase such as LipB transfers an octanoyl group from octanoyl-acyl carrier protein (ACP) to the target protein, after which lipoyl synthase LipA inserts sulfur atoms at C6 and C8 to generate the mature lipoyl group (Jordan & Cronan, 1997a, 1997b, 2003). More recently, an alternative pathway for protein lipoylation has been identified that is widely distributed across both bacterial and archaeal lineages (Cao, Koch, et al., 2018; Jin, Hachisuka et al., 2020; Jin, Sato, Hachisuka, & Atomi, 2022; Neti, Sil et al., 2022; Tanabe et al., 2023). We termed the lipoate:protein ligases involved sLpl(AB). Their archaeal origin distinguishes them evolutionarily from the previously characterized ligases (Tanabe et al., 2023). sLpl enzymes can initiate de novo protein lipoylation from free octanoate, after which lipoamide assembly is completed by the concerted action of the two unusual enzymes LipS1 and LipS2, which replace LipA and introduce the two sulfur atoms required to convert the octanoyl moiety into lipoamide. Although the LipS1/S2-type lipoyl synthase is less common than the canonical LipA-dependent pathway, it is remarkably widespread among prokaryotes (Jin et al., 2020; Jin et al., 2022; Neti et al., 2022; Tanabe et al., 2023).

Just as the classical salvage and de novo pathways for protein lipoylation coexist in many organisms, including *Escherichia coli* (Cronan, 2016, 2024), the sLpl–LipS1/S2 pathway may occur alongside one or both of these pathways. This raises fundamental questions about substrate ranges and the mechanisms by which parallel lipoylation pathways distinguish between their respective targets. Initial evidence showed that the GcvH-like LbpA proteins involved in sHdr-dependent sulfur oxidation are not recognized by the canonical lipoyl-attachment machineries of *E. coli* and *Bacillus subtilis* (Cao, Koch, et al., 2018). Moreover, *B. subtilis* octanoyltransferase LipM failed to modify LbpA acceptor proteins from bacterial sulfur oxidizers *in vitro*, and the sulfur oxidizer sLpl homologs failed to complement the growth of *E. coli lpl(AB)* and *lipB* deletion strains, suggesting that the ligases were specific for the cognate LbpA1 and LbpA2 acceptor proteins (Cao, Koch, et al., 2018). Conversely, sLpl(AB) lipoate:protein ligases from sulfur-oxidizing bacteria exhibited cross-species functionality among sulfur oxidizers, while failing to recognize lipoyl domains or proteins from organisms lacking components of an sHdr–LbpA sulfur-oxidation system (Cao, Koch, et al., 2018). However, the molecular basis of this substrate specificity has not yet been systematically investigated. This question is particularly relevant in sulfur-oxidizing bacteria ranging from the Alphaproteobacterium *Hyphomicrobium denitrificans* to lithoautotrophic Gammaproteobacteria and more distantly related members of the phylum Aquificota, which all encode the structurally and sequence-related but functionally distinct lipoylation substrates GcvH on the one hand and LbpA1 and LbpA2 on the other. How the lipoylation machineries discriminate between these homologous proteins to ensure their correct post-translational modification remains largely unclear. This study addresses whether the sLpl-LipS1/S2-based pathway is specifically dedicated to LbpA proteins, whether this specificity is already established at the level of substrate recognition by the ligases, and, if so, which molecular features determine this discrimination.

## 2. RESULTS

### 2.1 Setting the stage: choice of protein substrates and lipoate:protein ligases from model organisms

Bacteria encoding the sLpl-LipS1/S2 assembly pathway retain the canonical pathways for maturation of lipoamide (Cao, Koch, et al., 2018). The latter pathways appear restricted to the lipoate-binding components of 2-oxoacid dehydrogenases and glycine cleavage systems, whereas the distinct substrates of the novel pathway are specifically recognized by sLpl(AB) and subsequently modified by LipS1/LipS2. We hypothesized that substrate recognition and pathway discrimination occur at the initial octanoate/lipoate:protein ligase-catalyzed step. To test this hypothesis, we selected a representative set of model organisms based on the presence of the sLpl-LipS1/S2-dependent pathway, canonical lipoate biosynthesis enzymes, potential lipoylation substrates, and the sHdr sulfur oxidation pathway (Table 1).

**TABLE 1.**
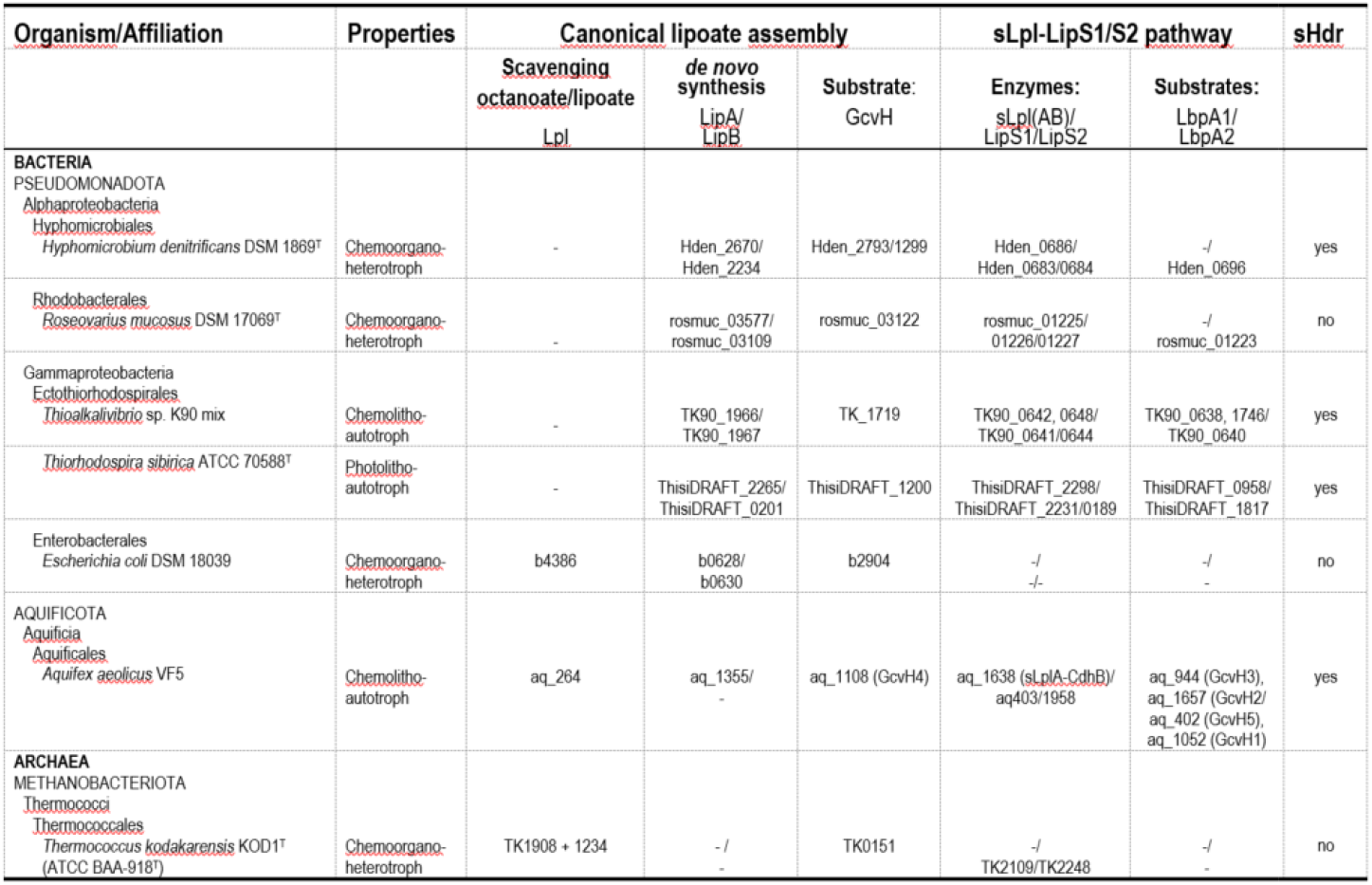
Occurrence of the canonical lipoate biosynthesis routes and the sLpl-LipS1/S2 pathway, their probable substrates and the sHdr pathway of sulfur oxidation in the model organisms of this study. The table also lists the archaeal organism *Thermococcus kodakarensis*, a hyperthermophilic chemoorganoeheterotroph incapable of dissimilatory sulfur oxidation, in which the LipS1/S2 lipoate synthases have been discovered (Jin et al., 2020). The five lipoylation target proteins from *Aquifex aeolicus* were previously tested for their ability to function as glycine cleavage protein H components in *B. subtilis* (Cao, Hong et al., 2018). The identifiers GcvH1 to GcvH5 assigned in that study are also provided in the present table for reference.

In the chosen model organisms, proteins representing potential lipoylation substrates and enzymes resembling lipoate:protein ligases were analyzed with respect to their genetic context (Fig. 2) and subjected to phylogenetic analyses (Fig. 3). This enabled the classification of lipoylation substrates as LbpA or GcvH proteins and the sequence-based distinction of Lpl from sLpl ligases.

**FIGURE 3.**
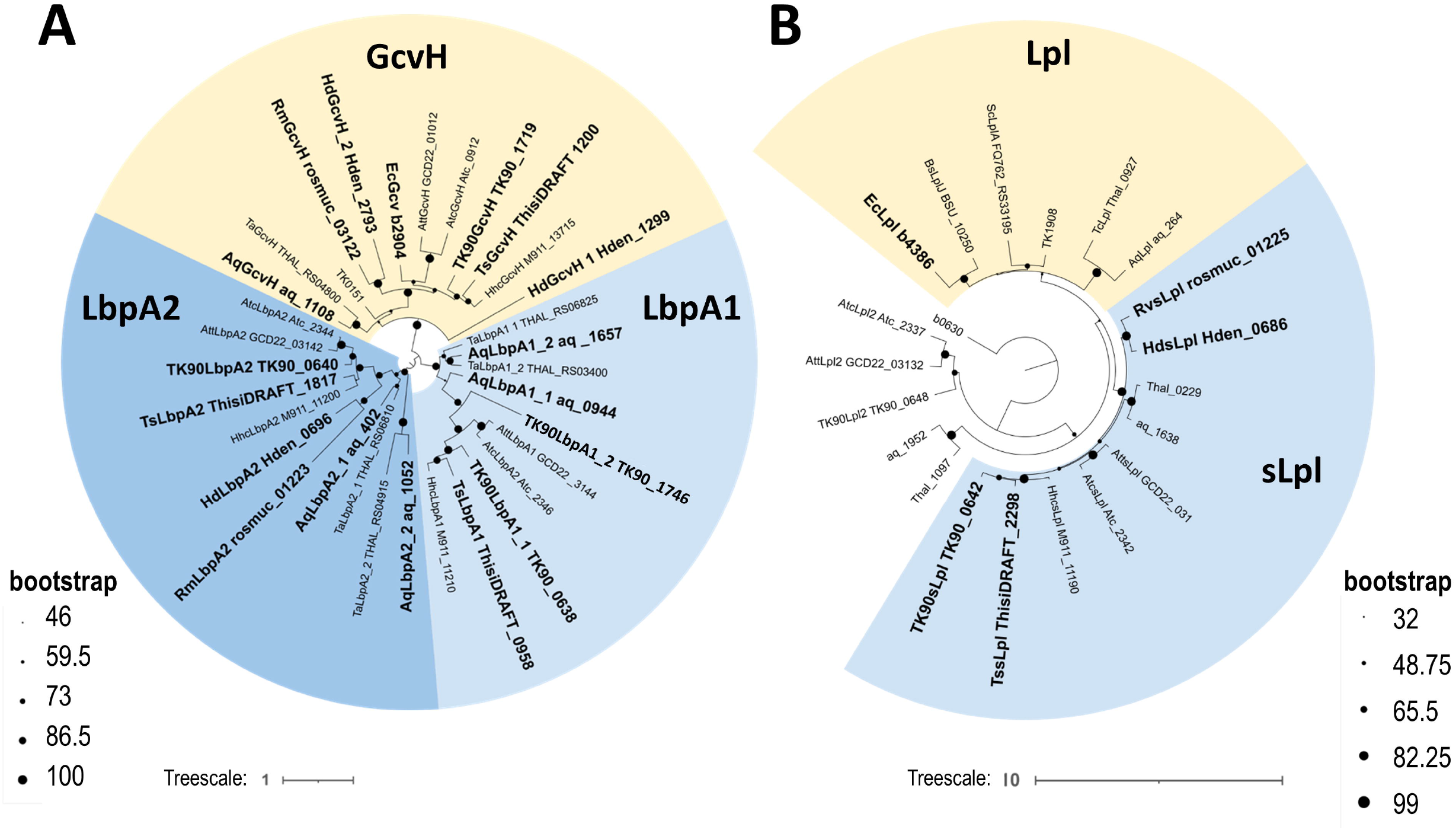
Phylogenetic trees for LbpA and GcvH proteins (A) and lipoate:protein ligases (B). The tree for the lipoate:protein ligases was rooted with the distantly related lipoate/octanoyl transferase LipB (b6030) from *E. coli* as an outgroup. The evolutionary history was inferred using the Maximum Likelihood method. Consensus trees are shown. Bootstrap values given at branching points are based on 3000 replicates. All positions containing gaps and missing data were eliminated. The treefiles are available as supplementary material. The 18 lipoylation substrates and the 5 lipoate:protein ligases studied in this work are highlighted in bold in Fig. 3A and 3B, respectively.

The chemoorganoheterotrophic Alphaproteobacterium *Hyphomicrobium denitrificans* DSM 1869^T^ is a well-established genetic model system for investigating the sLpl-LipS1/LipS2-based lipoate assembly pathway. In *H. denitrificans*, this pathway is necessary for maturation of the *Hd*LbpA2 protein, which operates together with the sHdr system for sulfur oxidation (Cao, Koch, et al., 2018; Tanabe et al., 2023). In addition, *H. denitrificans* is capable of *de novo* lipoate biosynthesis via LipB and LipA (Table 1).

Substrates for lipoylation include not only HdLbpA2 (Hden_0696) and *Hd*GcvH_2 (Hden_2793), which is a classical GcvH protein, but also the GcvH-like protein *Hd*GcvH_1 (Hden_1299) (Figs. 2,3), that probably serves as electron donor to OsmC (Hden_1301), a hydroperoxide reductase combatting reactive oxygen species (Cussiol et al., 2010; Cussiol, Alves, de Oliveira, & Netto, 2003; Lesniak, Barton, & Nikolov, 2003; Piccirillo, Alegria et al., 2018). *Roseovarius mucosus* DSMZ 17069^T^ (Biebl, Allgaier et al., 2005) is an example for an organism equipped with genes for the sLpl-LipS1/LipS2 pathway but incapable of sHdr-based sulfur oxidation (Table 1). The probable LbpA1-like substrate for lipoylation via sLpl-LipS1/S2 is encoded downstream of genes *fdhGBFD-fdsD* (Fig. 2) for cytoplasmic formate dehydrogenase and proteins involved in sulfuration of its bis-molybdopterin guanine dinucleotide cofactor (Hartmann & Leimkühler, 2013). LipB and LipA are present and probably responsible for lipoylation of *R. mucosus* GcvH (Table 1, Fig. 2). The chemolithoautotroph *Thioalkalivibio* sp. K90mix (Muyzer, Sorokin et al., 2011) and the photolithoautotroph *Thiorhodospira sibirica* ATCC 700588^T^ (Bryantseva, Gorlenko et al., 1999) are members of the family Ectothiorhodospiraceae within the Gammaproteobacteria. The organisms contain two to three sulfur oxidation-related LbpA proteins. All of these proteins, except TK90_1746, have previously been characterized as recombinant proteins (Cao, Koch, et al., 2018). The *Thioalkalivibrio* sp. K90mix sLpl(AB) TK90_0642 has been characterized as an octanoate/lipoate:protein ligase (Cao, Koch, et al., 2018). An obligately chemolithoautotrophic, hyperthermophilic member of the bacterial phylum Aquificota, *Aquifex aeolicus* VF5 (Deckert, Warren et al., 1998), is the fifth model organism. *A. aeolicus* harnesses energy from elemental sulfur or thiosulfate and contains the sulfur-oxidizing sHdr system including four different probable LbpA proteins (Fig. 2). In addition, *A. aeolicus* encodes one GcvH-like protein (Table 1, Figs. 2, 3). All five proteins have previously been tested for functionality as glycine cleavage protein H components and as substrates for octanoyl transferase (LipM) in *Bacillus subtilis* (Cao, Hong, et al., 2018).

### 2.2. Production and validation of Lpl and sLpl lipoate:protein ligases and their potential target proteins

When produced heterologously in *E. coli* BL21 (DE3) as recombinant proteins, lipoylation targets are predominantly obtained as non-lipoylated apoproteins due to the low endogenous concentration of EcLpl(AB), which is insufficient to catalyze lipoylation (Yang, Wang et al., 2025). This provided an ideal basis for testing a broad range of apoproteins as substrates for in vitro lipoylation. To this end, 18 different lipoylation targets (5 LbpA1, 6 LbpA2, 7 GcvH) from the selected model organisms were produced as recombinant proteins, together with five different lipoate:protein ligases (Fig. 2). Potential lipoate:protein ligases from *Aquifex aeolicus* were not included in our analyses because they originate from a hyperthermophilic organism and are expected to require elevated temperatures for optimal activity. Such temperatures would likely have compromised the stability of the substrate proteins, which were predominantly derived from mesophilic model organisms. The activity of each ligase was verified using its respective native substrate and also related target proteins. In addition, optimal reaction conditions for each enzyme were established by optimizing incubation time and temperature. It was also confirmed that all ligases accept octanoate as a substrate in addition to lipoate. Representative experiments are shown in Fig. S1.

Some *shdr-lbpA* gene clusters in sulfur-oxidizing bacteria, including those of *Acidithiobacillus caldus*, contain an additional *lpl*-like gene. *Thioalkalivibrio* sp. K90mix is another example (Fig. 3). Phylogenetic analysis (Fig. 2B) revealed that the corresponding proteins form a distinct clade, separate from both classical lipoate ligases, exemplified by the *E. coli* enzyme, and the sLpl enzymes. As a proof of principle, we produced the protein encoded by TK90_0648 from *Thioalkalivibrio* sp. K90mix in *E. coli* and tested its activity using recombinant apo-LbpA2 from the same organism as substrate. The enzyme was indeed capable of lipoylating apo-*TK90*LbpA2, although with very low efficiency. Complete lipoylation within 15 min required equimolar concentrations of enzyme and substrate (Fig. S2). The ligase was therefore not investigated any further in the context of the current study.

### 2.3 Testing lipoate:protein ligases for specificity

With a broad range of potential substrates and several highly active lipoate:protein ligases available, all possible substrate–ligase combinations were systematically tested. The resulting lipoylation reactions were analyzed by native gel-shift assays (Fig. 4), exploiting the difference in net charge between the apo- and holoproteins. Because lipoylation of a lysine residue neutralizes its positive charge, the holoproteins carry one net positive charge less than the corresponding apoproteins and therefore migrate faster towards the anode during native polyacrylamide gel electrophoresis. Except for reactions yielding unambiguous results, all observed mobility shifts were independently confirmed by mass spectrometry (Figs. S3-S18). Mass spectrometry was also the only suitable method for analyzing *TK90*LbpA1_1, which did not produce sufficiently well-resolved bands in native polyacrylamide gels.

**FIGURE 4.**
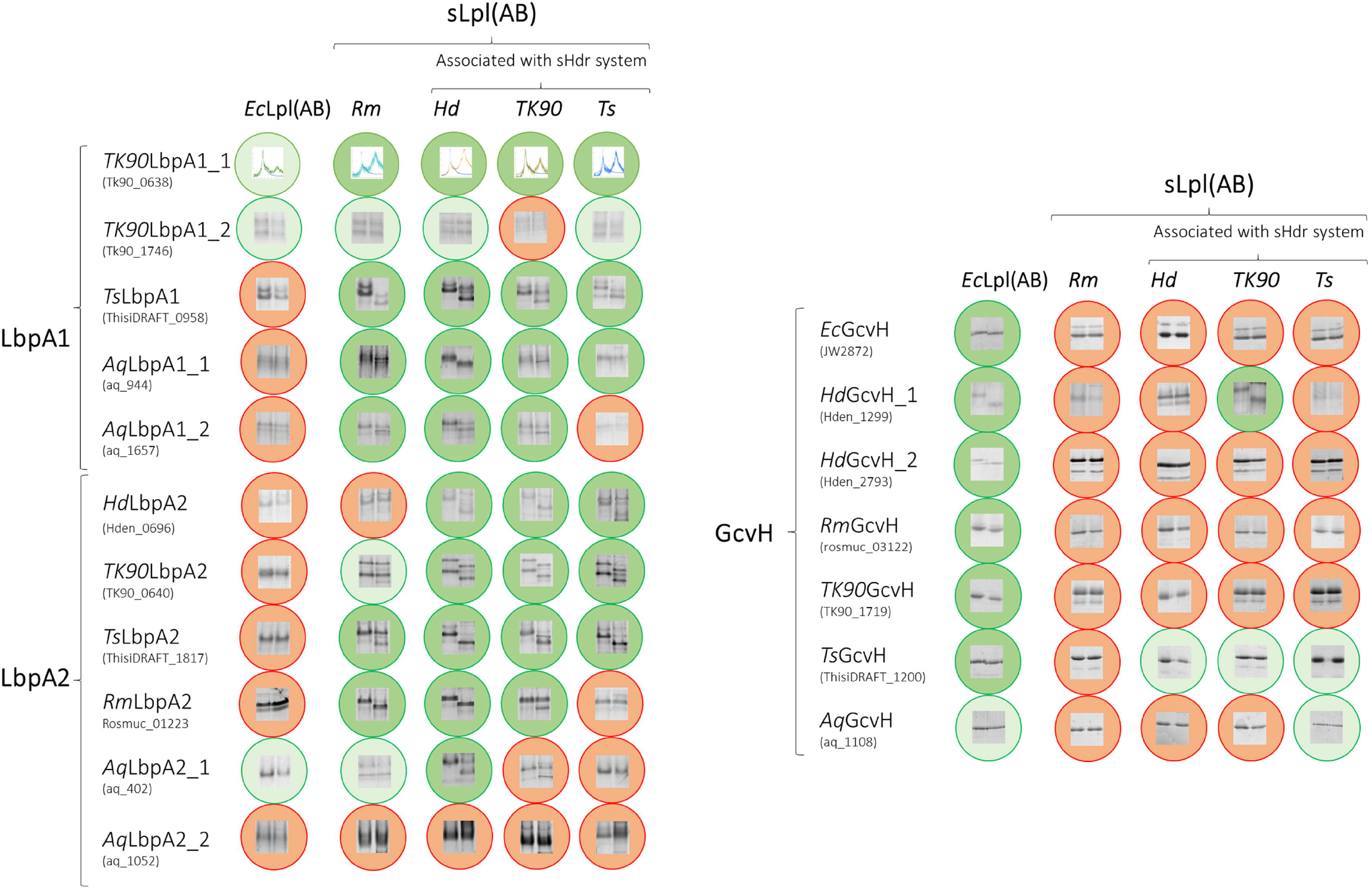
Summary of substrate recognition by lipoate:protein ligases. Substrate specificity was analyzed by native gel-shift assays. The concentration of acceptor proteins was 10 μM in all reactions, and the respective ligase was added at 0.5 μM. Positive reactions are indicated by a green background, whereas low-efficiency lipoylation detectable only by mass spectrometry is indicated by a light-green background. Negative results are indicated by red circles. *Ec, Escherichia coli*; *Rm, Roseovarius mucosus*; *Hd, Hyphomicrobium denitrificans*; *TK90, Thioalkalivibrio* sp. K90mix; *Ts, Thiorhodospira sibirica*; *Aq, Aquifex aeolicus*.

Lpl(AB) from *E. coli* has been reported to exhibit a broad substrate range and to catalyze the lipoylation of GcsH (glycine cleavage system protein H) from both algae (Yang et al., 2025) and human cells (Bick et al., 2024). Consistent with this broad substrate specificity, *Ec*Lpl(AB) recognized all bacterial GcvH proteins tested, irrespective of their organism of origin. In contrast, *Ec*Lpl(AB) failed to lipoylate the vast majority of the LbpA proteins tested, and those that were modified showed only very low levels of lipoylation (Fig. 4, Figs. S3–S18). The sLpl(AB) enzymes displayed an essentially opposite substrate preference: they efficiently modified the vast majority of LbpA proteins, irrespective of their organism of origin, but failed to recognize GcvH-like proteins, even when derived from the same organism (Fig. 4, Figs. S3–S18). *Aq*LbpA2_2 represented a notable exception, as it was not recognized by any of the ligases tested.

### 2.4 Protein surface charge as a determinant for recognition of lipoylation targets

An alignment of GcvH and LbpA proteins revealed a strictly conserved lysine residue that serves as the site of lipoylation in all proteins (Fig. 5 (Cao, Koch, et al., 2018)). With a few exceptions, including LbpA2_2 from *A. aeolicus*, the most conspicuous common feature of the LbpA proteins is the presence of two conserved cysteine residues that are absent from GcvH proteins and are located near the N- and C-termini, respectively (Fig. 5 (Cao, Koch, et al., 2018)). These conserved cysteines appeared to be plausible determinants of substrate recognition by sLpl(AB)-type lipoate ligases. To test this possibility, LbpA2 from *Thioalkalivibrio* sp. K90mix was produced with Cys-to-Ala substitutions at positions 7 and 145, individually and in combination, and the resulting variants were tested as substrates for the sLpl(AB) enzyme from the same organism. Gel-shift assays combined with mass spectrometric analysis unambiguously demonstrated efficient lipoylation of all three variants (Fig. S19). We therefore conclude that the conserved cysteine residues are not required for recognition of TK90LbpA2 by its cognate sLpl(AB) lipoate:protein ligase and are probably also dispensable for substrate recognition of LbpA proteins in general, although they may nevertheless have an important functional role.

**FIGURE 5.**
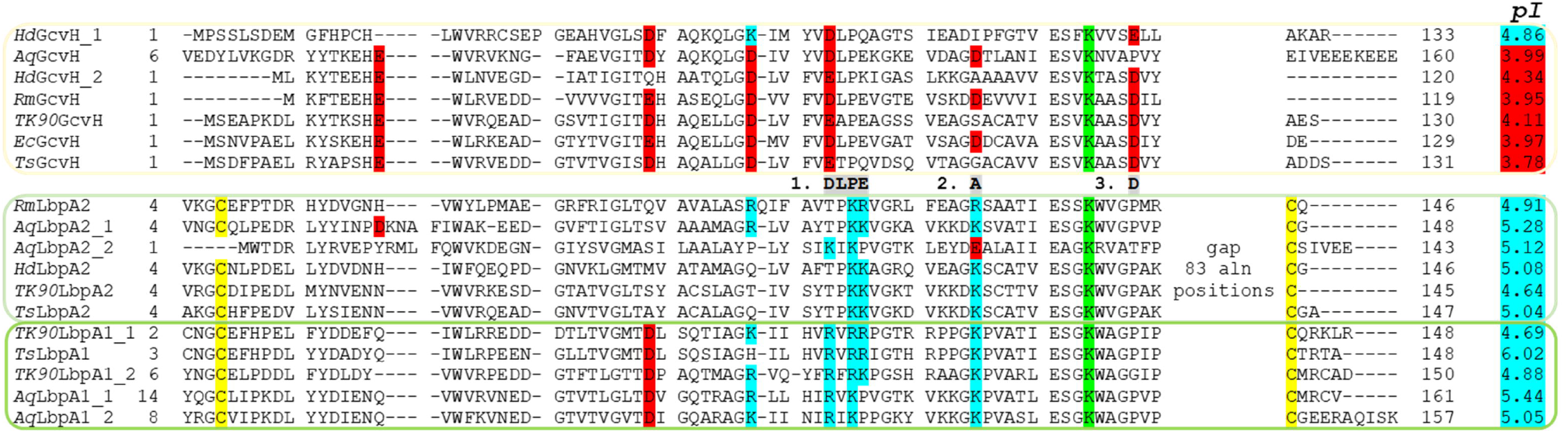
Partial sequence alignment of the 18 GcvH and LbpA proteins studied in this work. The conserved lysine residue binding the lipoyl moiety is highlighted in green. Conserved basic or acidic amino acid residues are highlighted in blue or red, respectively. The calculated isoelectric point (*pI*) is given for each protein. Values below 4.5 are colored red, those above are highlighted in light blue. The line above the *Roseovarius mucosus* protein *Rm*LbpA2 indicates the amino acid substitutions introduced at the corresponding positions in variants 1, 2, and 3.

In addition to the conserved cysteine residues, LbpA and GcvH proteins exhibit characteristic differences in the distribution of charged amino acids. GcvH proteins share several conserved acidic residues that are frequently replaced at corresponding positions in LbpA proteins by neutral or, more commonly, basic residues such as lysine and arginine (Fig. 5). These differences are reflected not only in the generally lower *pI* values of GcvH proteins (Fig. 5) but also in pronounced differences in their local surface charge. Accordingly, the surface of *E. coli* GcvH that interacts with its cognate lipoate ligase, as defined by the crystal structure of the LplA–GcvH complex (Fujiwara et al., 2010), is predominantly negatively charged (Fig. 6). In contrast, the corresponding surface of *R. mucosus* LbpA2, shown as a representative example in Fig. 6A, is predominantly positively charged. This distinction extends to all proteins examined in this study: acidic residues are consistently enriched on the corresponding surface of GcvH proteins, whereas such an enrichment is absent from both LbpA1 and LbpA2 proteins (Fig. S20).

**FIGURE 6.**
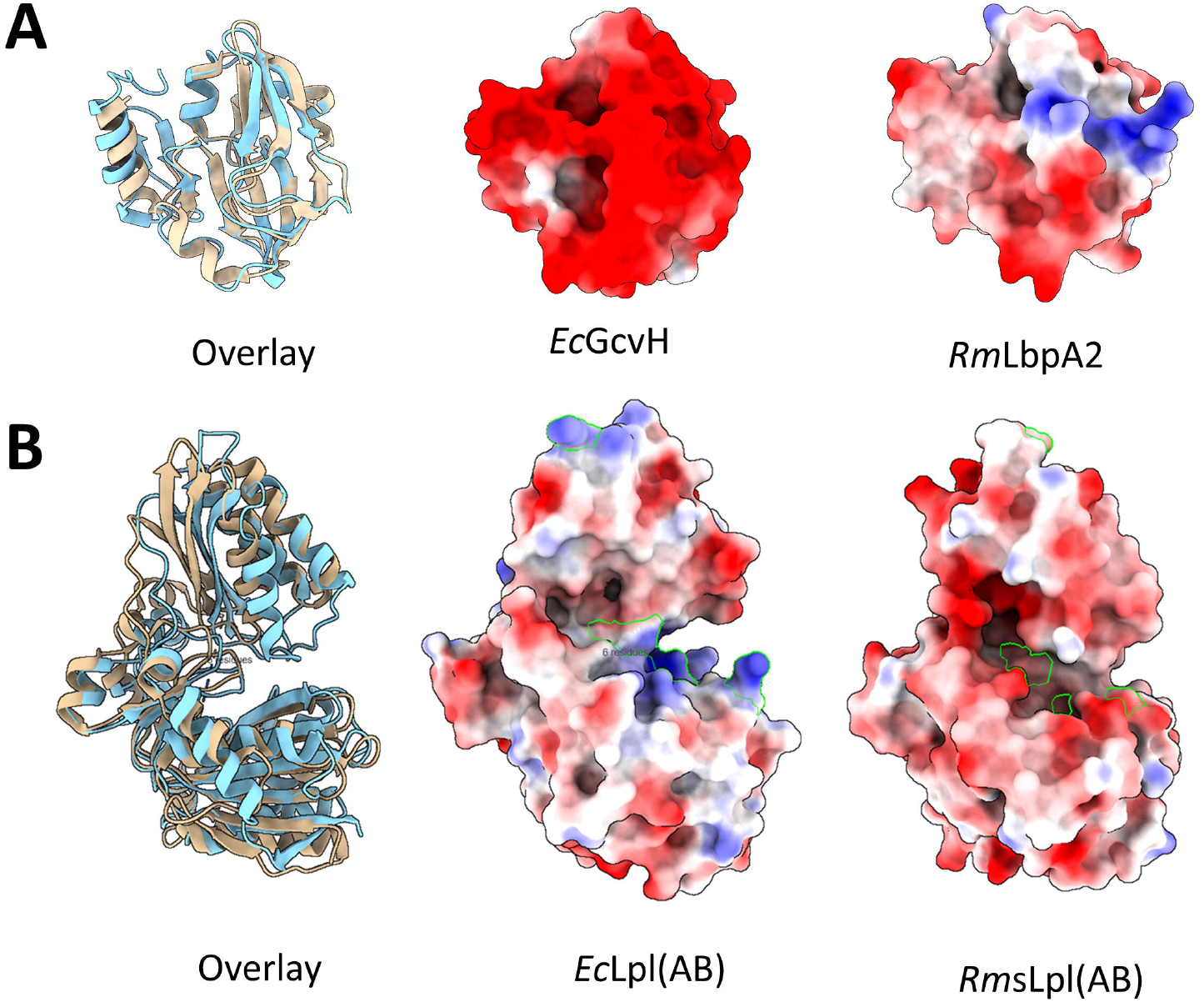
Structural comparison of selected bacterial lipoylation targets and lipoate:protein ligases. **(A)** Comparison of *E. coli* GcvH (beige, (PDB 3AB9 (Okamura-Ikeda et al., 2010)) and LbpA2 (rosmuc_01223) from *Roseovarius mucosus*, modelled by AlphaFold (Jumper, Evans et al., 2021; Varadi, Bertoni et al., 2024). The left panel provides an overlay of the *E. coli* (beige) and *R. mucosus* (blue) proteins. The middle and right panels show the structures of *E. coli* GcvH and *R. mucosus* LbpA2 in the same orientation and colored according to surface electrostatic potential, respectively. Red, acidic; blue, basic; white, neutral. **(B)** Comparison of *E. coli* Lpl(AB) (1×2G (Fujiwara, Toma et al., 2005) and sLpl(AB) from *R. mucosus* predicted by AlphaFold (Jumper et al., 2021; Varadi et al., 2024). The left panel provides an overlay of the *E. coli* (beige) and *R. mucosus* (blue) proteins. The middle and right panels show the structures of *E. coli* Lpl(AB) and *R. mucosus* sLpl(AB) in the same orientation and colored according to surface electrostatic potential, respectively. Red, acidic; blue, basic; white, neutral. The basic amino acid residues of the *E. coli* ligase, Arg^47^, Arg^140^, Lys^143^, Lys^175^, and Lys^272^, that contribute to the hydrogen-bonding interactions with apo-*Ec*GcvH (Fujiwara et al., 2010), are highlighted by green outlines. The corresponding positions in the *Roseovarius* ligase are marked in the same way, and are occupied by Leu^50^, Phe^147^, Gly^151^, His^182^, and Gly^283^, that are predominantly uncharged at neutral pH.

To experimentally assess the contribution of these sequence and surface-charge differences to substrate recognition, we selected LbpA2 from *R. mucosus* (*Rm*LbpA2) as a representative LbpA protein for systematic mutagenesis. Amino acid substitutions were introduced at three positions indicated in Fig. 5, either individually or in combination. The resulting recombinant variant proteins were produced in and purified from *E. coli* and tested as substrates for *Rm*sLpl(AB) and *E. coli* Lpl(AB) using native PAGE mobility-shift assays (Fig. 7).

**FIGURE 7.**
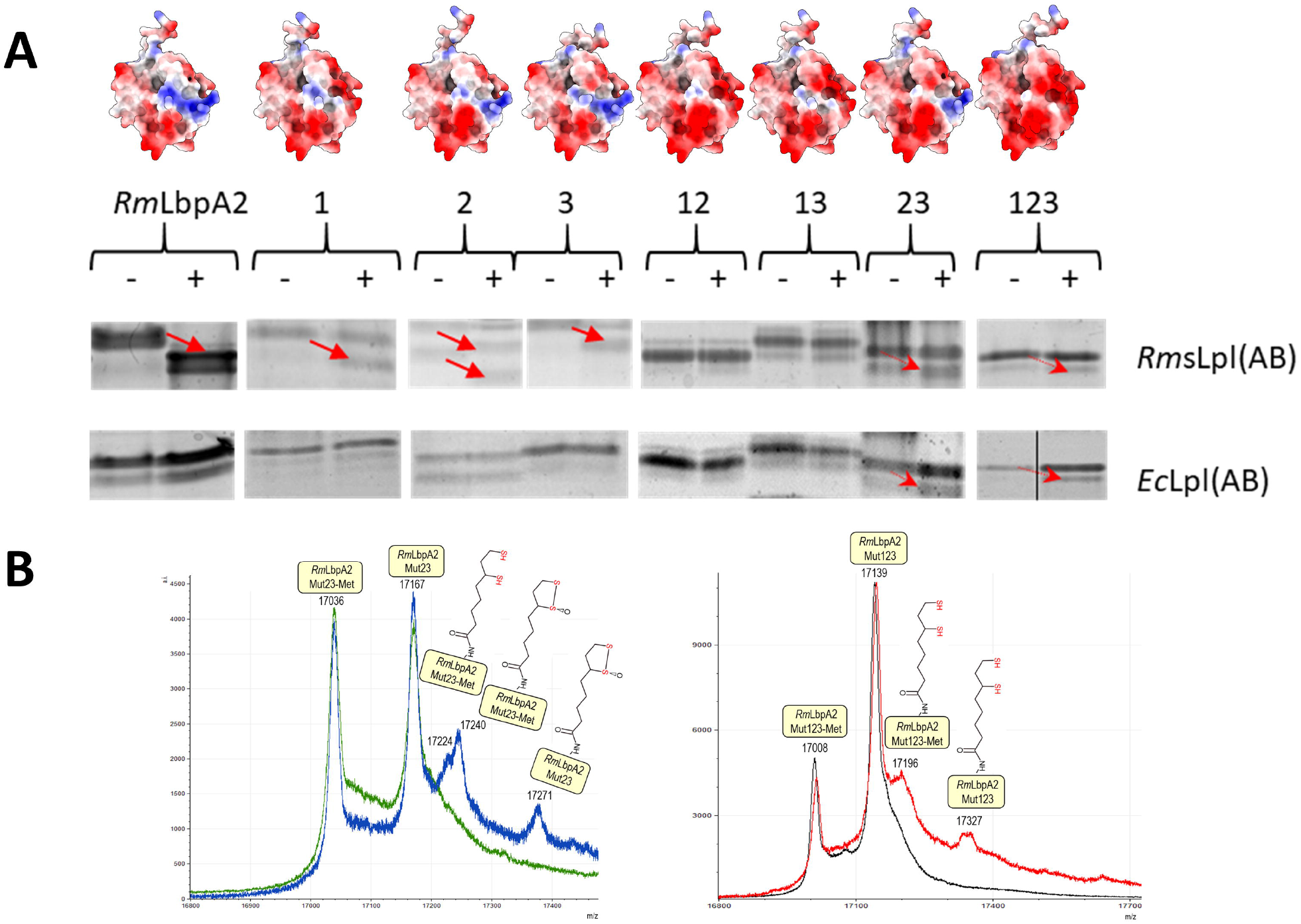
In vitro post-translational modification of wild-type and variant RmLbpA2 protein. (A) Recombinant Strep-tagged LbpA2 from *Roseovarius mucosus* and variants carrying changes at positions 1, 2, or 3 (indicated in Fig. 5), either individually or in combinations (12, 13, 23 or 123) were incubated with sLpl(AB) from the same organism (*Rm*sLpl(AB)) or with Lpl(AB) from *E. coli* and subsequently analyzed by native PAGE. Reactions contained 10 μM lipoylation target and 0.5 μM of the respective ligase and were incubated for 45 min at 30 ºC or 37ºC for the enzymes from *R. mucosus* and *E. coli*, respectively. Red arrows indicate a shift from the apoprotein to the faster-migrating holoprotein, demonstrating complete or partial lipoylation. Structural models of the protein variants are shown above the gels. The structures were predicted using AlphaFold (Jumper et al., 2021; Varadi et al., 2024) and colored according to surface electrostatic potential (red, acidic; blue, basic; white, neutral). All proteins are shown in the same orientation. (B) Mass spectrometric verification of lipoylation of *Rm*LbpA2 carrying substitutions at positions 2 and 3 (*Rm*LbpA_Mut23) or at all three positions (*Rm*LbpA_Mut123) by *Ec*Lpl(AB). For the *Rm*LbpA2_Mut23 variant, *in vitro* lipoylation resulted in a mass increase of 205 Da, consistent with attachment of a mono-oxygenated lipoyl moiety (+189 Da + 16 Da) corresponding to the thiosulfinate form of lipoate, a known oxidation product (Krishnan & Garnett, 2011; Reed, DeBusk, Gunsalus, & Hornberger Jr., 1951; Reed, Gunsalus et al., 1953; Stary, Jindal, & Murray, 1975). Recombinant *Rm*LbpA2 was purified from *E. coli* as a mixture of polypeptides retaining or lacking the initiator methionine.

As observed previously, wild-type *Rm*LbpA2 was efficiently and completely lipoylated by its cognate *Rm*sLpl(AB), whereas it was not recognized by *Ec*Lpl(AB) (Fig. 7). Individual alteration of any one of the three selected positions (TPRK^51-54^DLPE, R^63^A or P^76^D) did not abolish recognition by *Rm*sLpl(AB), although lipoylation was no longer complete within the chosen reaction time of 45 min. None of these individual substitutions was sufficient to enable detectable lipoylation by *Ec*Lpl(AB). The picture changed when substitutions at positions 2 and 3, or at all three positions, were combined, resulting in a pronounced shift in the surface-charge distribution of *Rm*LbpA2 (Fig. 7A). These variants became substrates for *Ec*Lpl(AB) and underwent partial lipoylation, which was independently verified by mass spectrometry (Fig. 7B). Although the substitutions did not result in complete conversion of *Rm*LbpA2 into an *Ec*Lpl(AB) substrate, these findings provide strong evidence that electrostatic complementarity contributes substantially to discrimination between GcvH and LbpA proteins by the two types of lipoate:protein ligases. This interpretation is further supported by the complementary surface-charge properties of the ligases themselves: the substrate-binding surface of sLpl(AB) is predominantly negatively charged, whereas the corresponding region of the *E. coli* enzyme is predominantly positively charged (Fig. 6B).

## 3. DISCUSSION

Post-translational modification of specific lysine residues is essential for the function of GcvH, LbpA and other lipoyl domain-containing proteins. The present study provides insight into the molecular basis of substrate specificity at the level of the lipoate:protein ligases catalyzing the initial step in the attachment of lipoate or octanoate. Selective substrate recognition is particularly relevant in organisms such as *Thioalkalivibrio* spp. and *Thiorhodospira sibirica*, in which canonical and sLpl-LipS1/S2-driven lipoylation systems coexist and target GcvH as well as lipoyl domains of the pyruvate and 2-oxoglutarate dehydrogenase complexes alongside several sulfur oxidation-related LbpA proteins.

Extending earlier work (Cao, Koch, et al., 2018), the systematic *in vitro* analysis performed here revealed a clear separation in substrate specificity: *Ec*Lpl(AB) efficiently modified a range of GcvH proteins but showed little or no activity toward LbpA proteins, whereas sLpl(AB) enzymes strongly preferred LbpA proteins. Importantly, this preference was not restricted to cognate enzyme–substrate pairs but extended across species boundaries. Thus, recognition is determined by conserved properties that distinguish the GcvH and LbpA protein families rather than by species-specific interactions. The sLpl(AB) enzymes were furthermore shown to attach not only lipoate but also free octanoate to their acceptor proteins, confirming that, like canonical Lpl(AB) enzymes, they function as both lipoate and octanoate ligases (Cao, Koch, et al., 2018; Green, Morris et al., 1995; Morris, Reed, & Cronan, 1994, 1995; Zhao, Miller et al., 2003)

The high structural similarity between GcvH and LbpA proteins raises the question of which molecular features allow the ligases to discriminate between them. Unlike several other post-translational modifications for which target-site selection is strongly influenced by short linear sequence motifs surrounding the modified residue, as exemplified by protein phosphorylation and small ubiquitin-like modifier conjugation (SUMOylation) (Mok, Kim et al., 2010; Sampson, Wang, & Matunis, 2001), lipoylation does not appear to be primarily determined by the immediate sequence environment of the target lysine. Mutations of conserved residues flanking the lipoyl-lysine are well tolerated, indicating that the immediate sequence environment is not the primary determinant of ligase recognition (Wallis & Perham, 1994). Instead, natural lipoylation substrates share a compact three-dimensional fold of approximately 80 residues suggesting that substrate recognition involves features of the folded acceptor protein (Reche & Perham, 1999). The two highly conserved cysteine residues located toward the N- and C-terminal regions of LbpA proteins initially appeared to represent one prominent feature distinguishing GcvH and LbpA proteins. In structural models, these residues are positioned sufficiently close to permit formation of an intramolecular disulfide bond in the oxidized state (Sun et al., 2017) (Fig. 1). However, replacement of either cysteine individually or of both residues simultaneously by alanine did not alter recognition by sLpl(AB). The conserved cysteine pair is therefore excluded as a primary determinant of LbpA recognition.

Another pronounced distinguishing feature emerged from comparison of the electrostatic surfaces of the two acceptor-protein families. Fujiwara et al. (2010) showed in the *Ec*Lpl(AB)–apo-*Ec*GcvH complex that recognition involves an extensive network of hydrogen-bonding interactions. Many of the participating residues in *Ec*GcvH are acidic, whereas several residues on the interacting surface of *Ec*Lpl(AB) are basic. These include Asp^43^, Glu^46^, Glu^61^, Asp^68^, and Asp^86^ in *Ec*GcvH (Fig. 5) and Arg^47^, Arg^140^, Lys^143^, Lys^175^, and Lys^272^ in *Ec*Lpl(AB) (Fig. 6B) (Fujiwara et al., 2010). The resulting charge complementarity led Fujiwara et al. to propose that electrostatic interactions are an important determinant of acceptor-protein specificity. Our comparisons strongly support this concept and extend it to the discrimination between GcvH and LbpA proteins. A positively charged region surrounding the relevant interaction surface of *Ec*Lpl(AB) is readily apparent (Fig. 6A) and complements the negatively charged surface of *Ec*GcvH (Fig. 6B). In sLpl(AB) enzymes, however, several of the basic residues occupying equivalent positions in *Ec*Lpl(AB) are replaced by residues that are neutral or oppositely charged (Fig. 6B). Conversely, LbpA proteins display a characteristic positively charged surface region where GcvH proteins are predominantly negatively charged (Fig. 6A). Thus, the electrostatic properties of both interaction partners appear to have evolved in a coordinated manner: canonical Lpl(AB) and GcvH form one complementary pair, whereas sLpl(AB) and LbpA form another. The *Rm*LbpA2 mutagenesis experiments provide direct experimental support for this interpretation. Replacement of individual regions containing basic residues did not identify a single amino acid that alone determined ligase specificity. Instead, recognition progressively depended on the combined properties of several surface regions. Alteration of two of the selected regions was already sufficient to severely impair efficient recognition by *Rm*sLpl(AB), demonstrating that the native distribution of surface charge is critical for productive interaction with the cognate ligase. Conversely, acquisition of negatively charged residues was sufficient to turn *Rm*LbpA2 into a substrate for *Ec*Lpl(AB). Detectable lipoylation, however, occurred only when two or three investigated regions had been altered simultaneously and even these variants were not completely modified. Thus, the substitutions were sufficient to shift substrate preference but not to fully convert *Rm*LbpA2 fully into a GcvH-like substrate.

Our observations argue against a simple recognition motif consisting of one or a few conserved residues. Rather, they support a model in which substrate selection is governed by a broader interaction surface whose electrostatic character results from the combined contribution of multiple residues. In this model, the three-dimensional fold provides the common structural framework that positions the lipoyl-lysine appropriately for catalysis, whereas the electrostatic properties of spatially separated surface regions determine which ligase can productively dock onto a particular acceptor protein.

Reports that lipoate:protein ligases can act on short synthetic peptides (Jin et al., 2022; Puthenveetil, Liu et al., 2009) do not necessarily contradict this model. Selected natural or engineered ligases can modify peptide substrates, and engineered *E. coli* Lpl(AB) variants have been exploited extensively for site-specific modification of short acceptor peptides (Fernandez-Suarez, Baruah et al., 2007; Uttamapinant, White et al., 2010). However, such peptide substrates are highly selected sequences and do not imply that the physiological discrimination between closely related natural acceptor proteins is governed by a short linear motif. The strong substrate preferences observed here across a large panel of intact GcvH and LbpA proteins, together with the effects of altering several spatially separated regions of *Rm*LbpA2, instead indicate that the folded protein surface is a major determinant of physiological substrate selection.

Taken together, our results suggest that GcvH and LbpA specificity is determined not by individual recognition residues but by broader surface properties, particularly electrostatic complementarity with their respective ligases. Further mutagenesis of both acceptor protein classes will help define these interaction surfaces and establish whether reciprocal changes in surface charge can reverse ligase specificity.

## 4 MATERIALS AND METHODS

### 4.1 Bacterial strains, plasmids, primers, and growth conditions

Table S1 lists the bacterial strains, and plasmids that were used for this study. Escherichia coli strains were grown on complex lysogeny broth (LB) medium (Bertani, 2004) under aerobic conditions at 37°C unless otherwise indicated. Escherichia coli strains 10-beta or DH5α were used for molecular cloning. E. coli BL21 (DE3) was used for recombinant protein production. Antibiotics for E. coli were used at the following concentrations (in μg ml^-1^): ampicillin, 100; kanamycin, 50.

### 4.2 Recombinant DNA techniques

Standard techniques for DNA manipulation and cloning were used unless otherwise indicated (Ausubel, Brent et al., 1997). Restriction enzymes, T4 ligase and Q5 polymerase were obtained from New England Biolabs (Ipswich, UK) and used according to the manusfacturer’s instructions. Oligonucleotides for cloning were obtained from Eurofins MWG (Ebersberg, Germany) and are listed in Table S2. Plasmid DNA from E. coli was purified using the GenJET Plasmid Miniprep Kit (Thermo Scientific, Waltham, USA). Chromosomal DNA from E. coli, Hyphomicrobium denitrificans, Thioalkalivibrio sp. K90mix, Rosevarius mucosus and Thiorhodospira sibirica strains was prepared using the Simplex easy DNA kit (GEN-IAL GmbH, Troisdorf, Germany).

### 4.3 Construction of plasmids for recombinant production of lipoate:protein ligases and LbpA/GcvH proteins

Aquifex aeolicus VF5 lbpA and gcvH genes were commercially synthesized and cloned into in pET28a(+) vectors (BioCat GmbH Heidelberg, Germany). The other genes for lipoate:protein ligases and their substrates were amplified by PCR with the primers listed in Table S2 and cloned into either pET22b(+) or pET28a(+) resulting in the plasmids listed in Table S1. Cysteine to alanine exchanges were implemented to TK90LbpA2 by SOE PCR using the primers listed in Table S2. Amino acid exchanges at positions 1, 2, and 3 in RmLbpA2 (Figure 5) were also implemented by SOE PCR with suitable primers (Table S2). Genes for RmLbpA2 variants with exchanges at positions 1 & 2 or 2 & 3 or 1 & 3 or 1, 2 & 3 were codon-optimized for E. coli and purchased from BioCat, Heidelberg, Germany. All recombinant proteins were produced in E. coli BL21 (DE3). Overnight precultures were used to inoculate fresh LB medium with a ratio of 1:50 (v/v). Synthesis of recombinant proteins was induced by the addition of 0.1 mM IPTG when cultures had reached an OD_600_ of 0.4–0.6, followed by incubation over night at 16ºC. Cells were harvested by centrifugation (11,000 × g, 20 min, 4ºC). Strep-tagged proteins were resuspended in 100 mM Tris–HCl buffer (pH 8.0) containing 150 mM NaCl and 2 mM dithiothreitol. His-tagged proteins were resuspended in buffer containing 50 mM sodium-phosphate, 300 mM NaCl, 2 mM dithiothreitol and 20 mM imidazole (pH 7.5). In all cases, lysozyme, RNase, DNase and EDTA-free Halt™ protease inhibitor cocktail (Thermo Fisher Scientific, Waltham, USA) were added according to the manufacturers’ instructions. Cells were lysed by sonication and insoluble cell material was subsequently removed by ultracentrifugation (145,000 × g, 60 min, 4ºC). His-tagged and Strep-tagged proteins were purified with Ni-NTA Superflow (IBA Lifesciences, Göttingen, Germany) and Strep-Tactin Superflow (IBA Lifesciences, Göttingen, Germany) or with StrepTrap™ columns (Cytiva, Marlborough, USA) using an ÄKTApurifier 900 (Cytiva, Marlborough, USA), respectively, according to the manufacturer’s instructions. All buffers contained 2 mM dithiothreitol. For storage, the proteins were transferred to 50 mM Tris–HCl buffer (pH 7.5) containing 100 mM NaCl and 1 mM dithiotreitol. Protein concentrations were determined with the Pierce BCA protein assay kit (Thermo Scientific, Dreieich, Germany). Pure recombinant proteins were quantified on the basis of their calculated extinction coefficients at 280 nm. Purity of protein preparations was assessed by sodium dodecyl sulfate-polyacrylamid gel electrophoresis (SDS-PAGE).

### 4.4 In vitro lipoylation assays

Lipoate ligase activity was assayed by observing a mobility shift of lipoate-binding proteins upon modification by native gel electrophoresis as originally described by Miles and Guest (Miles and Guest, 1987). In vitro lipoylation assays contained in a final volume of 200 μl, 50 mM Tri-Hcl (pH 7.5), 0.5 mM DTT, 1.5 mM α-lipoic acid or sodium octanoate, 1.5 mM MgCl_2_, 1.5 mM ATP, 10 μM lipoate-binding protein (LbpA or GcvH) and 0.5-1 μM of the respective ligase. Unless noted otherwise, the reaction mixtures (200 μl) were incubated at the optimum growth temperature of the source organism of the respective ligase (37ºC for E. coli, 34ºC for Thioalkalivibrio sp. K90 mix (Muyzer et al., 2011), 30ºC for T. sibirica (Bryantseva et al., 1999), 30ºC for H. denitrificans and 31ºC for R. mucosus (Biebl et al., 2005)) for 45 min and analyzed by native PAGE and mass spectrometry. Results were compared for assays with and without lipoic acid or octanoate and visualized via Native PAGE. GcvH and LbpA proteins were analyzed in 20% and 15% gels, respectively.

### 4.5 Mass spectrometry

Before MALDI-TOF mass spectrometry analysis, the samples were desalted and concentrated with ZipTip C18 (Merck Millipore, Burlington, USA) using 0.1% trifluoroacetic acid (TFA) as desalting solution and 70% acetonitrile/0.1% TFA as elution solution. One microliter of each sample was mixed with 1 μL of matrix α-cyano-4-hydroxycinnamic acid and analyzed by the Core Facility “Protein Synthesis and Bioanalytics” of the University of Bonn.

### 4.6 Phylogenetic tree inference

For inference of a phylogenetic trees for the lipoate:protein ligases and the LbpA and GcvH proteins, proteins were aligned using MAFFT (Katoh & Standley, 2013) and the alignments were trimmed by stripping columns with gaps. A maximum likelihood phylogeny was inferred using IQ-TREE (Nguyen, Schmidt, von Haeseler, & Minh, 2015; Trifinopoulos, Nguyen, von Haeseler, & Minh, 2016). The best amino acid substitution model was calculated with ModelFinder (Kalyaanamoorthy, Minh et al., 2017). In both cases, the LG+I+G4 model had the lowest BIC (Bayesian Information Criterion) score and was considered best. Branch support was calculated by ultrafast bootstrap (3000 replicates) (Hoang, Chernomor et al., 2018). Finally, the trees were displayed using iTol (Letunic & Bork, 2021).

## Supporting information

Treefile LbpA-GcvH proteins

Treefile Lipoate:protein ligases

## SUPPLEMENTARY MATERIAL

Additional supporting information may be found online in the supporting Information section at the end of this article

## Author contributions

C.K., C.D. designed the experiments. C.K., E.H., T.S.T. and M.G.M. performed the experiments. C.K., T.S.T., M.G.M. and C.D. analyzed the data. C.K. and C.D. wrote the manuscript, C.D.: funding acquisition, project administration, supervision.

## ACKNOWLEDGEMENTS

This work was funded by the Deutsche Forschungsgemeinschaft (Grant Da 351/14-1). We gratefully acknowledge the support of the Core Facility “Protein Synthesis and Bioanalytics” of the University of Bonn.

## REFERENCES

Abramson, J., Adler, J., Dunger, J., Evans, R., Green, T., Pritzel, A., Ronneberger, O., Willmore, L., Ballard, A. J., Bambrick, J., Bodenstein, S. W., Evans, D. A., Hung, C. C., O’Neill, M., Reiman, D., Tunyasuvunakool, K., Wu, Z., Zemgulyte, A., Arvaniti, E., Beattie, C., Bertolli, O., Bridgland, A., Cherepanov, A., Congreve, M., Cowen-Rivers, A. I., Cowie, A., Figurnov, M., Fuchs, F. B., Gladman, H., Jain, R., Khan, Y. A., Low, C. M. R., Perlin, K., Potapenko, A., Savy, P., Singh, S., Stecula, A., Thillaisundaram, A., Tong, C., Yakneen, S., Zhong, E. D., Zielinski, M., Zidek, A., Bapst, V., Kohli, P., Jaderberg, M., Hassabis, D., & Jumper, J. M. (2024). Accurate structure prediction of biomolecular interactions with AlphaFold 3. Nature, 630(8016), 493–500.

Acosta, I. C., Albers, A., Fang, L., Serrato, G., Teoh, W. P., Glanville, D. G., & Alonzo, F., 3rd. (2025). Staphylococcus aureus exploits lipoic acid salvage to combat host oxidative stress. Cell Reports, 44(8), 116095.

Ausubel, F. A., Brent, R., Kingston, R. E., Moore, D. D., Seidman, J. G., Smith, J. A., & Struhl, K. (1997). Current protocols in molecular biology. New York: John Wiley & Sons.

Bertani, G. (2004). Lysogeny at mid-twentieth century: P1, P2, and other experimental systems. Journal of Bacteriology, 186(3), 595–600.

Biebl, H., Allgaier, M., Lunsdorf, H., Pukall, R., Tindall, B. J., & Wagner-Dobler, I. (2005). Roseovarius mucosus sp. nov., a member of the Roseobacter clade with trace amounts of bacteriochlorophyll a. International Journal of Systematic and Evolutionary Microbiology, 55(Pt 6), 2377–2383.

Brocklehurst, S. M., & Perham, R. N. (1993). Prediction of the three-dimensional structures of the biotinylated domain from yeast pyruvate carboxylase and of the lipoylated H-protein from the pea leaf glycine cleavage system: a new automated method for the prediction of protein tertiary structure. Protein Sci., 2(4), 626–639.

Bryantseva, I. A., Gorlenko, V. M., Kompantseva, E. I., Imhoff, J. F., Sling, J., & Mityushina, L. (1999). Thiorhodospira sibirica gen. nov., sp. nov., a new alkaliphilic purple sulfur bacterium from a Siberian soda lake. International Journal of Systematic Bacteriology, 49, 697–703.

Cao, X., Hong, Y., Zhu, L., Hu, Y., & Cronan, J. E. (2018). Development and retention of a primordial moonlighting pathway of protein modification in the absence of selection presents a puzzle. Proceedings of the National Academy of Sciences of the United States of America, 115(4), 647–655.

Cao, X., Koch, T., Steffens, L., Finkensieper, J., Zigann, R., Cronan, J. E., & Dahl, C. (2018). Lipoate-binding proteins and specific lipoate-protein ligases in microbial sulfur oxidation reveal an atypical role for an old cofactor. Elife, 7, e37439.

Cronan, J. E. (2016). Assembly of lipoic acid on its cognate enzymes: an extraordinary and essential biosynthetic pathway. Microbiology and Molecular Biology Reviews, 80(2), 429–450.

Cronan, J. E. (2024). Lipoic acid attachment to proteins: stimulating new developments. Microbiology and Molecular Biology Reviews, 88(2), e0000524.

Cronan, J. E., Zhao, X., & Jiang, Y. (2005). Function, attachment and synthesis of lipoic acid in Escherichia coli. Advances in Microbial Physiology, 50, 103–146.

Cussiol, J. R., Alegria, T. G., Szweda, L. I., & Netto, L. E. (2010). Ohr (organic hydroperoxide resistance protein) possesses a previously undescribed activity, lipoyl-dependent peroxidase. Journal of Biological Chemistry, 285(29), 21943–21950.

Cussiol, J. R., Alves, S. V., de Oliveira, M. A., & Netto, L. E. (2003). Organic hydroperoxide resistance gene encodes a thiol-dependent peroxidase. Journal of Biological Chemistry, 278(13), 11570–11578.

Deckert, G., Warren, P. V., Gaasterland, T., Young, W. G., Lenox, A. L., Graham, D. E., Overbeek, R., Snead, M. A., Keller, M., Aujay, M., Huber, R., Feldman, R. A., Short, J. M., Olsen, G. J., & Swanson, R. V. (1998). The complete genome of the hyperthermophilic bacterium Aquifex aeolicus. Nature, 392(6674), 353–358.

Fernandez-Suarez, M., Baruah, H., Martinez-Hernandez, L., Xie, K. T., Baskin, J. M., Bertozzi, C. R., & Ting, A. Y. (2007). Redirecting lipoic acid ligase for cell surface protein labeling with small-molecule probes. Nature Biotechnology, 25(12), 1483–1487.

Fujiwara, K., Maita, N., Hosaka, H., Okamura-Ikeda, K., Nakagawa, A., & Taniguchi, H. (2010). Global conformational change associated with the two-step reaction catalyzed by Escherichia coli lipoate-protein ligase A. Journal of Biological Chemistry, 285(13), 9971–9980.

Fujiwara, K., Toma, S., Okamura-Ikeda, K., Motokawa, Y., Nakagawa, A., & Taniguchi, H. (2005). Crystal structure of lipoate-protein ligase A from Escherichia coli. Determination of the lipoic acid-binding site. Journal of Biological Chemistry, 280(39), 33645–33651.

Green, D. E., Morris, T. W., Green, J., Cronan, J. E., Jr., & Guest, J. R. (1995). Purification and properties of the lipoate protein ligase of Escherichia coli. Biochemical Journal, 309 (Pt 3), 853–862.

Hartmann, T., & Leimkühler, S. (2013). The oxygen-tolerant and NAD+-dependent formate dehydrogenase from Rhodobacter capsulatus is able to catalyze the reduction of CO2 to formate. FEBS J, 280(23), 6083–6096.

Hoang, D. T., Chernomor, O., von Haeseler, A., Minh, B. Q., & Vinh, L. S. (2018). UFBoot2: Improving the ultrafast bootstrap approximation. Molecular Biology and Evolution, 35(2), 518–522.

Jin, J. Q., Hachisuka, S. I., Sato, T., Fujiwara, T., & Atomi, H. (2020). A structurally novel lipoyl synthase in the hyperthermophilic archaeon Thermococcus kodakarensis. Applied and Environmental Microbiology, 86(23), e01359–01320.

Jin, J. Q., Sato, T., Hachisuka, S. I., & Atomi, H. (2022). A lipoate-protein ligase is required for de novo lipoyl-protein biosynthesis in the hyperthermophilic archaeon Thermococcus kodakarensis. Applied and Environmental Microbiology, 88(13), e0064422.

Jones, D. D., Horne, H. J., Reche, P. A., & Perham, R. N. (2000). Structural determinants of post-translational modification and catalytic specificity for the lipoyl domains of the pyruvate dehydrogenase multienzyme complex of Escherichia coli. Journal of Molecular Biology, 295(2), 289–306.

Jones, D. D., Stott, K. M., Howard, M. J., & Perham, R. N. (2000). Restricted motion of the lipoyl-lysine swinging arm in the pyruvate dehydrogenase complex of Escherichia coli. Biochemistry, 39(29), 8448–8459.

Jordan, S. W., & Cronan, J. E., Jr. (1997a). Biosynthesis of lipoic acid and posttranslational modification with lipoic acid in Escherichia coli. Methods in Enzymology, 279, 176–183.

Jordan, S. W., & Cronan, J.E., Jr. (1997b). A new metabolic link. The acyl carrier protein of lipid synthesis donates lipoic acid to the pyruvate dehydrogenase complex in Escherichia coli and mitochondria. Journal of Biological Chemistry, 272(29), 17903–17906.

Jordan, S. W., & Cronan, J.E., Jr. (2003). The Escherichia coli lipB gene encodes lipoyl (octanoyl)-acyl carrier protein:protein transferase. Journal of Bacteriology, 185(5), 1582–1589.

Jumper, J., Evans, R., Pritzel, A., Green, T., Figurnov, M., Ronneberger, O., Tunyasuvunakool, K., Bates, R., Zidek, A., Potapenko, A., Bridgland, A., Meyer, C., Kohl, S. A. A., Ballard, A. J., Cowie, A., Romera-Paredes, B., Nikolov, S., Jain, R., Adler, J., Back, T., Petersen, S., Reiman, D., Clancy, E., Zielinski, M., Steinegger, M., Pacholska, M., Berghammer, T., Bodenstein, S., Silver, D., Vinyals, O., Senior, A. W., Kavukcuoglu, K., Kohli, P., & Hassabis, D. (2021). Highly accurate protein structure prediction with AlphaFold. Nature, 596(7873), 583–589.

Kalyaanamoorthy, S., Minh, B. Q., Wong, T. K. F., von Haeseler, A., & Jermiin, L. S. (2017). ModelFinder: fast model selection for accurate phylogenetic estimates. Nature Methods, 14(6), 587–589.

Katoh, K., & Standley, D. M. (2013). MAFFT multiple sequence alignment software version 7: improvements in performance and usability. Molecular Biology and Evolution, 30(4), 772–780.

Kikuchi, G., Motokawa, Y., Yoshida, T., & Hiraga, K. (2008). Glycine cleavage system: reaction mechanism, physiological significance, and hyperglycinemia. Proceedings of the Japan Acacemy of Sciences, Series B Physical and Biological Sciences, 84(7), 246–263.

Koch, T., & Dahl, C. (2018). A novel bacterial sulfur oxidation pathway provides a new link between the cycles of organic and inorganic sulfur compounds. ISME Journal, 12(10), 2479–2491.

Krishnan, C. V., & Garnett, M. (2011). Electrochemical behavior of the super antioxidant, α-lipoic acid. International Journal of Electrochemical Science, 6(8), 3607–3630.

Kümpel, C., Grosser, M., Tanabe, T. S., & Dahl, C. (2024). Fe/S proteins in microbial sulfur oxidation. Biochimica et Biophysica Acta (BBA) - Molecular Cell Research, 1871, 119732.

Lesniak, J., Barton, W. A., & Nikolov, D. B. (2003). Structural and functional features of the Escherichia coli hydroperoxide resistance protein OsmC. Protein Science, 12(12), 2838–2843.

Letunic, I., & Bork, P. (2021). Interactive Tree Of Life (iTOL) v5: an online tool for phylogenetic tree display and annotation. Nucleic Acids Research, 49(W1), W293-W296.

Lin, C. H., Chin, Y., Zhou, M., Sobol, R. W., Hung, M. C., & Tan, M. (2024). Protein lipoylation: mitochondria, cuproptosis, and beyond. Trends in Biochemical Sciences, 49(8), 729–744.

Mohr, M. G., Tanabe, T. S., Grosser, M., & Dahl, C. (2026). Biochemistry and transcriptional regulation of bacterial sulfur oxidation. Advances in Microbial Physiology, 89, 51–154.

Mok, J., Kim, P. M., Lam, H. Y., Piccirillo, S., Zhou, X., Jeschke, G. R., Sheridan, D. L., Parker, S. A., Desai, V., Jwa, M., Cameroni, E., Niu, H., Good, M., Remenyi, A., Ma, J. L., Sheu, Y. J., Sassi, H. E., Sopko, R., Chan, C. S., De Virgilio, C., Hollingsworth, N. M., Lim, W. A., Stern, D. F., Stillman, B., Andrews, B. J., Gerstein, M. B., Snyder, M., & Turk, B. E. (2010). Deciphering protein kinase specificity through large-scale analysis of yeast phosphorylation site motifs. Science Signaling, 3(109), ra12.

Morris, T. W., Reed, K. E., & Cronan, J.E., Jr. (1994). Identification of the gene encoding lipoate-protein ligase A of Escherichia coli. Molecular cloning and characterization of the lplA gene and gene product. Journal of Biological Chemistry, 269(23), 16091–16100.

Morris, T. W., Reed, K. E., & Cronan, J.E., Jr. (1995). Lipoic acid metabolism in Escherichia coli: the lplA and lipB genes define redundant pathways for ligation of lipoyl groups to apoprotein. Journal of Bacteriology, 177(1), 1–10.

Muyzer, G., Sorokin, D. Y., Mavromatis, K., Lapidus, A., Foster, B., Sun, H., Ivanova, N., Pati, A., D’Haeseleer, P., Woyke, T., & Kyrpides, N. C. (2011). Complete genome sequence of Thioalkalivibrio sp. K90mix. Standards in Genomic Sciences, 5(3), doi:10.4056/sigs.2315092.

Neti, S. S., Sil, D., Warui, D. M., Esakova, O. A., Solinski, A. E., Serrano, D. A., Krebs, C., & Booker, S. J. (2022). Characterization of LipS1 and LipS2 from Thermococcus kodakarensis: Proteins annotated as biotin synthases, which together catalyze formation of the lipoyl cofactor. ACS Bio Med Chem Au, 2(5), 509–520.

Nguyen, L. T., Schmidt, H. A., von Haeseler, A., & Minh, B. Q. (2015). IQ-TREE: a fast and effective stochastic algorithm for estimating maximum-likelihood phylogenies. Molecular Biology and Evolution, 32(1), 268–274.

Nyvltova, E., Smutna, T., Tachezy, J., & Hrdy, I. (2016). OsmC and incomplete glycine decarboxylase complex mediate reductive detoxification of peroxides in hydrogenosomes of Trichomonas vaginalis. Molecular and Biochemical Parasitology, 206(1-2), 29-38.

Okamura-Ikeda, K., Hosaka, H., Maita, N., Fujiwara, K., Yoshizawa, A. C., Nakagawa, A., & Taniguchi, H. (2010). Crystal structure of aminomethyltransferase in complex with dihydrolipoyl-H-protein of the glycine cleavage system: implications for recognition of lipoyl protein substrate, disease-related mutations, and reaction mechanism. Journal of Biological Chemistry, 285(24), 18684–18692.

Pares, S., Cohen-Addad, C., Sieker, L., Neuburger, M., & Douce, R. (1994). X-ray structure determination at 2.6-A resolution of a lipoate-containing protein: the H-protein of the glycine decarboxylase complex from pea leaves. Proceedings of the National Academy of Sciences of the United States of America, 91(11), 4850–4853.

Perham, R. N. (2000). Swinging arms and swinging domains in multifunctional enzymes: catalytic machines for multistep reactions. Annual Review of Biochemistry, 69, 961–1004.

Piccirillo, E., Alegria, T. G. P., Discola, K. F., Cussiol, J. R. R., Domingos, R. M., de Oliveira, M. A., Rezende, L., Netto, L. E. S., & Amaral, A. T. (2018). Structural insights on the efficient catalysis of hydroperoxide reduction by Ohr: Crystallographic and molecular dynamics approaches. PLoS One, 13(5), e0196918.

Puthenveetil, S., Liu, D. S., White, K. A., Thompson, S., & Ting, A. Y. (2009). Yeast display evolution of a kinetically efficient 13-amino acid substrate for lipoic acid ligase. Journal of the American Chemical Society, 131(45), 16430–16338.

Reche, P., & Perham, R. N. (1999). Structure and selectivity in post-translational modification: attaching the biotinyl-lysine and lipoyl-lysine swinging arms in multifunctional enzymes. EMBO Journal, 18(10), 2673–2682.

Reed, L. J., DeBusk, B. G., Gunsalus, I. C., & Hornberger Jr., C.S. (1951). Crystalline α-lipoic acid: a catalytic agent associated with pyruvate dehydrogenase. Science, 114(2952), 93–94.

Reed, L. J., Gunsalus, I. C., Schnakenberg, G. H. F., Soper, Q. F., Boaz, H. E., Kern, S. F., & Parke, T. V. (1953). Isolation, characterization and structure of α-lipoic acid. Journal of the American Chemical Society, 75(6), 1267–1270.

Rowland, E. A., Snowden, C. K., & Cristea, I. M. (2018). Protein lipoylation: an evolutionarily conserved metabolic regulator of health and disease. Current Opinion in Chemical Biology, 42, 76–85.

Sampson, D. A., Wang, M., & Matunis, M. J. (2001). The small ubiquitin-like modifier-1 (SUMO-1) consensus sequence mediates Ubc9 binding and is essential for SUMO-1 modification. Journal of Biological Chemistry, 276(24), 21664–21669.

Spalding, M. D., & Prigge, S. T. (2010). Lipoic acid metabolism in microbial pathogens. Microbiology and Molecular Biology Reviews, 74(2), 200–228.

Stary, F. E., Jindal, S. J., & Murray, R. W. (1975). Oxidation of α-lipoic acid. Journal of Organic Chemistry, 40(1), 58–62.

Sun, M. A., Wang, Y., Zhang, Q., Xia, Y., Ge, W., & Guo, D. (2017). Prediction of reversible disulfide based on features from local structural signatures. BMC Genomics, 18(1), 279.

Tanabe, T. S., Grosser, M., Hahn, L., Kümpel, C., Hartenfels, H., Vtulkin, E., Flegler, W., & Dahl, C. (2023). Identification of a novel lipoic acid biosynthesis pathway reveals the complex evolution of lipoate assembly in prokaryotes. PLoS Biology, 21(6), e3002177.

Trifinopoulos, J., Nguyen, L. T., von Haeseler, A., & Minh, B. Q. (2016). W-IQ-TREE: a fast online phylogenetic tool for maximum likelihood analysis. Nucleic Acids Research, 44(W1), W232–235.

Uttamapinant, C., White, K. A., Baruah, H., Thompson, S., Fernandez-Suarez, M., Puthenveetil, S., & Ting, A. Y. (2010). A fluorophore ligase for site-specific protein labeling inside living cells. Proceedings of the National Academy of Sciences of the United States of America, 107(24), 10914–10919.

Varadi, M., Bertoni, D., Magana, P., Paramval, U., Pidruchna, I., Radhakrishnan, M., Tsenkov, M., Nair, S., Mirdita, M., Yeo, J., Kovalevskiy, O., Tunyasuvunakool, K., Laydon, A., Zidek, A., Tomlinson, H., Hariharan, D., Abrahamson, J., Green, T., Jumper, J., Birney, E., Steinegger, M., Hassabis, D., & Velankar, S. (2024). AlphaFold Protein Structure Database in 2024: providing structure coverage for over 214 million protein sequences. Nucleic Acids Research, 52(D1), D368–D375.

Wallis, N. G., & Perham, R. N. (1994). Structural dependence of post-translational modification and reductive acetylation of the lipoyl domain of the pyruvate dehydrogenase multienzyme complex. Journal of Molecular Biology, 236(1), 209–216.

Yang, R., Wang, Y., Kong, M., Hu, Z., Zhang, Z., Shen, K., Meng, J., & Zeng, A. P. (2025). Boosting energy metabolism and biosynthesis in diverse organisms by a common bacterial salvage lipoylation protein. Nature Communications, 16(1), 7540.

Zhao, X., Miller, J. R., Jiang, Y., Marletta, M. A., & Cronan, J. E. (2003). Assembly of the covalent linkage between lipoic acid and its cognate enzymes. Chemical Biology, 10(12), 1293–1302.

