## Supplementary Figures and tables for "Substrate specificity in lipoylation pathways: How lipoate:protein ligases discriminate between protein substrates"

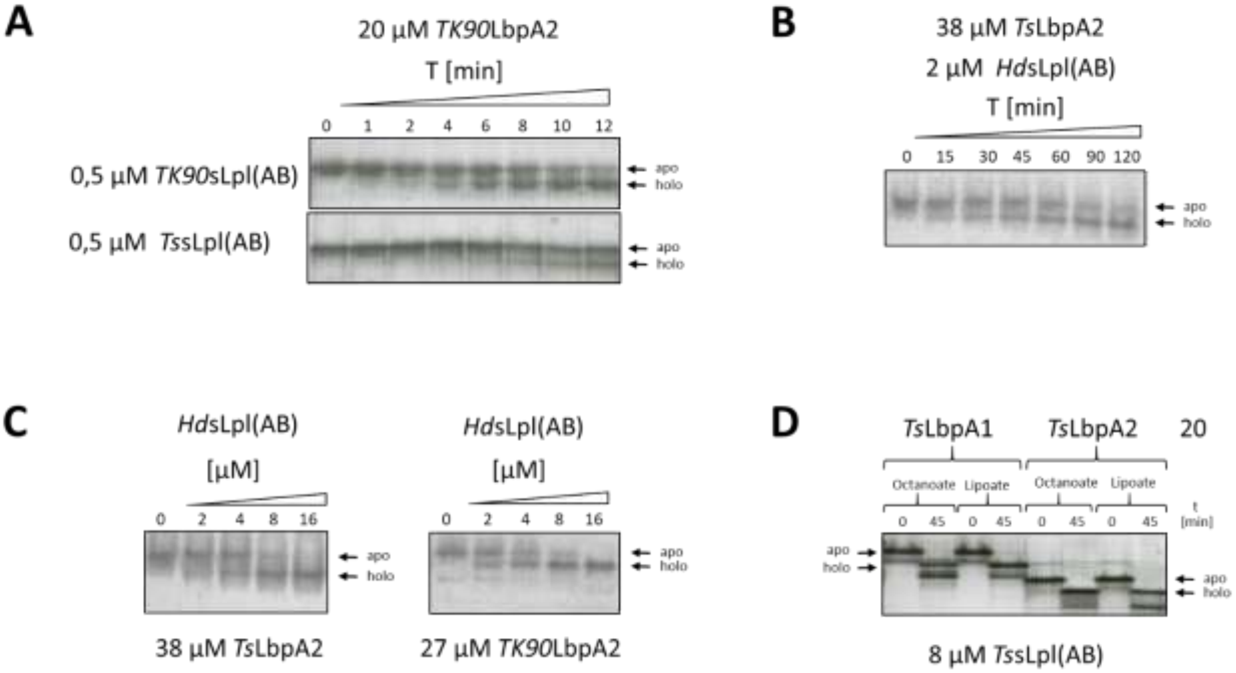


**Fig. S1. Representative activity assays for lipoate:protein ligases. (A)** sLpl(AB) from *Thioalkalivibrio* sp. K90mix (*TK90*sLpl(AB)) and *Thiorhdodspira sibirica* (*Ts*sLpl(AB)) catalyze lipoylation of LbpA2 from *Thioalkalivibrio* sp. K90mix (*TK90*LbpA2) in a time dependent manner at 30°C and 34°C, respectively. **(B)** sLpl(AB) from *Hyphomicrobium denitrificans* (*Hd*sLpl(AB)) catalyzes lipoylation of LbpA2 from *T. sibirica* (*Ts*LbpA2) in a time dependent manner. Reactions were run at 30°C. **(C)** Lipoylation efficiency of *Ts*LbpA2 and *TK90*LbpA2 increases with increasing concentration of *Hd*sLpl(AB). Reactions were run at 30°C for 30 min. **(D)** *Ts*sLpl(AB) catalyzes both octanyolation and lipoylation of *Ts*LbpA1 and *Ts*LbpA2 in the presence of 1.5 mM sodium octanoate or 1.5 mM lipoic acid. Reactions were run at 30°C for 45 min and mass shifts were verified by mass spectrometry.

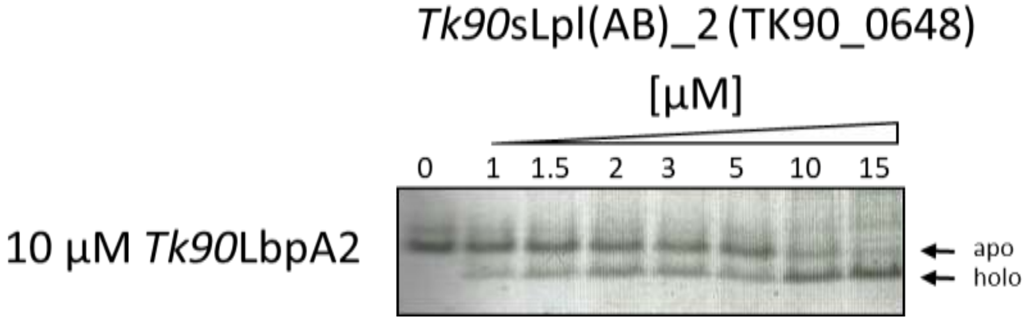


**Fig. S2. The enzyme encoded by TK90_0648 (TK90_Lpl(AB)2) is active as lipoate:protein ligase.** The ligase was produced as a recombinant protein with a carboxy-terminal His-tag. Reactions were incubated at 34$^{o}$C for 45 min. Note that complete lipoylation of 10 µM TK90_LbpA2 required 15 µM of the ligase, while 0.5 µM TK90sLpl(AB) encoded by TK90_0642 led to almost complete lipoylation of 20 µM of the same protein substrate within only 12 min.

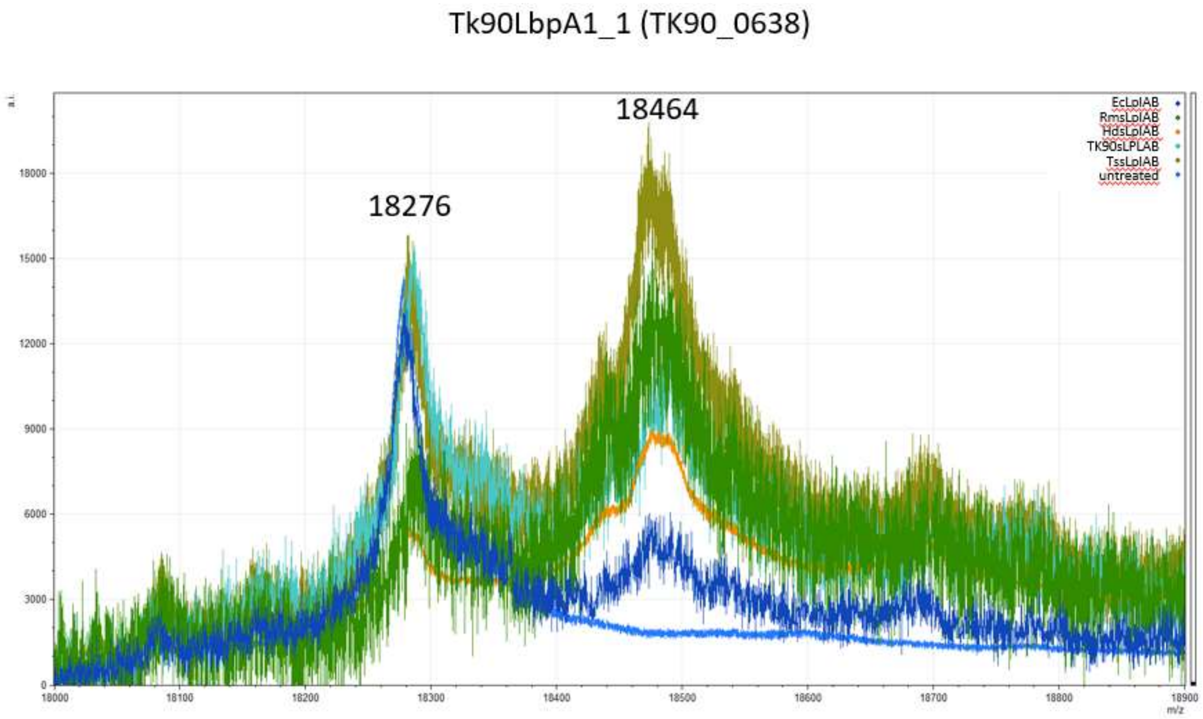


**Fig. S3. Mass spectrometry of *TK90*LbpA1_1 after incubation with different lipoate:protein ligases in the presence of 1.5 mM α-lipoic acid.** Efficient lipoylation was catalyzed by all sLpl(AB)s, whereas very little lipoylation was observed in the presence of *Ec*Lpl(AB).

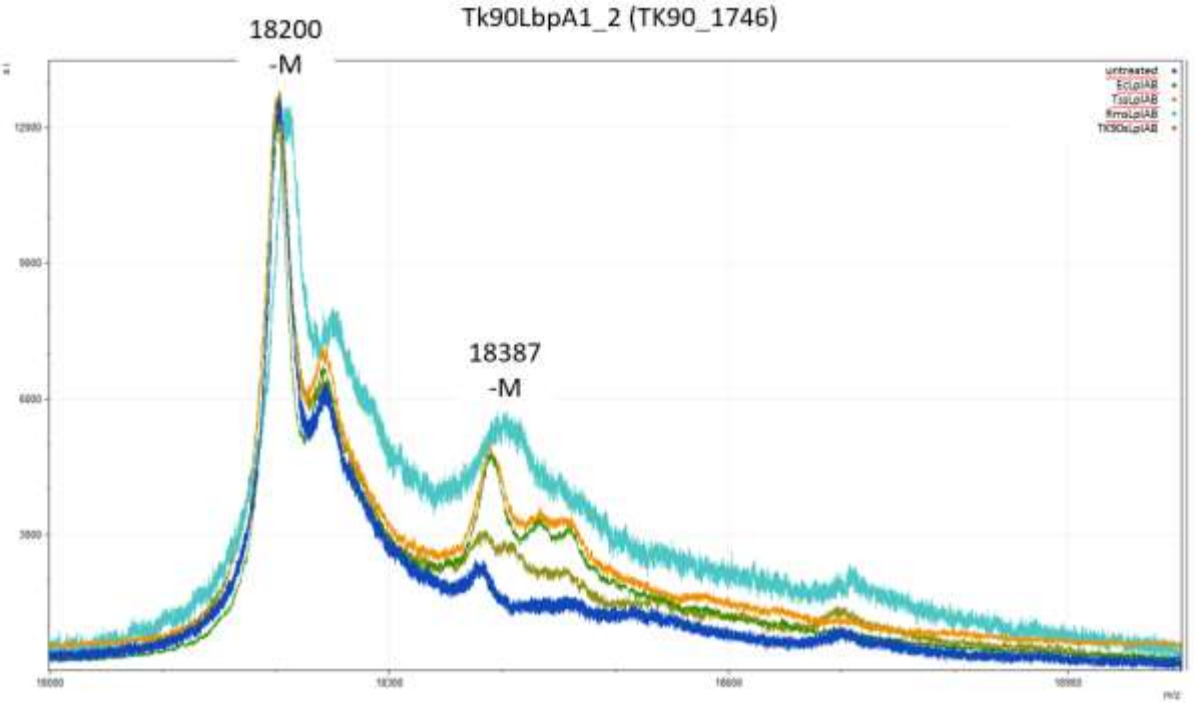


**Fig. S4. Mass spectrometry of *TK90*LbpA1_2 after incubation with different lipoate:protein ligases in the presence of 1.5 mM α-lipoic acid.** A small extent of lipoylation is achieved with *E. coli* Lpl(AB), *R. mucosus* sLpl(AB) and *T. sibirica* sLpl(AB). –M, the detected mass corresponds to the full-length polypeptide lacking the initiator methionine.

TsLbpA1 (ThisiDRAFT_0958)

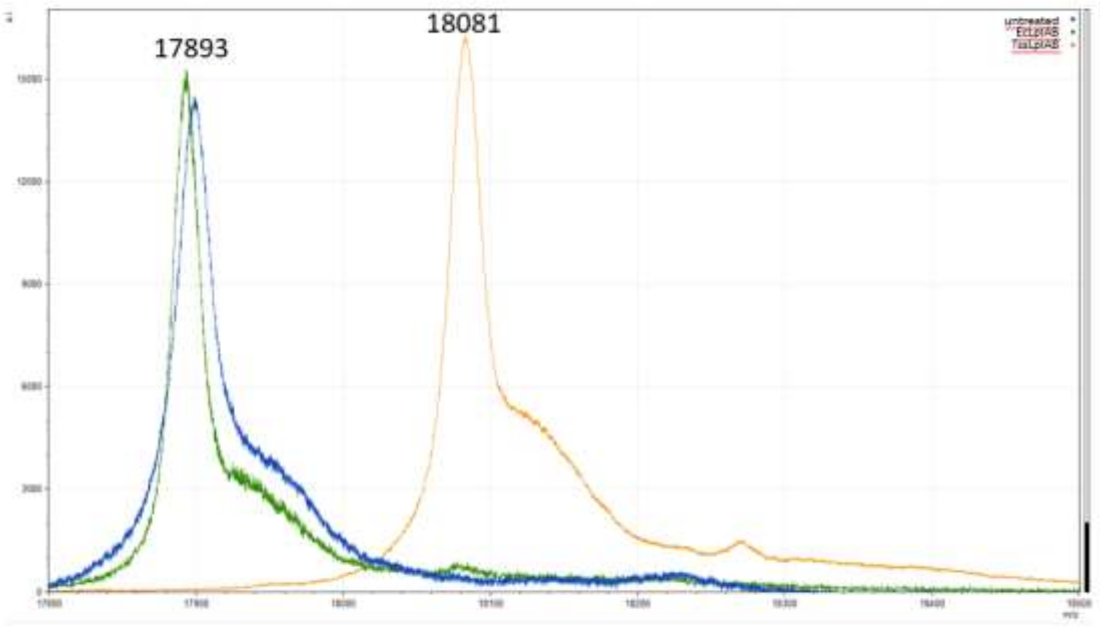


**Fig. S5. Mass spectrometry of *Ts*LbpA1 after incubation with different lipoate:protein ligases in the presence of 1.5 mM α-lipoic acid.** Efficient lipoylation was catalyzed sLpl(AB) from *Thiorhodospira sibirica,* whereas no modification was observed in the presence of *Ec*Lpl(AB).

AqLbpA1_1 (aq_944, GcvH3)

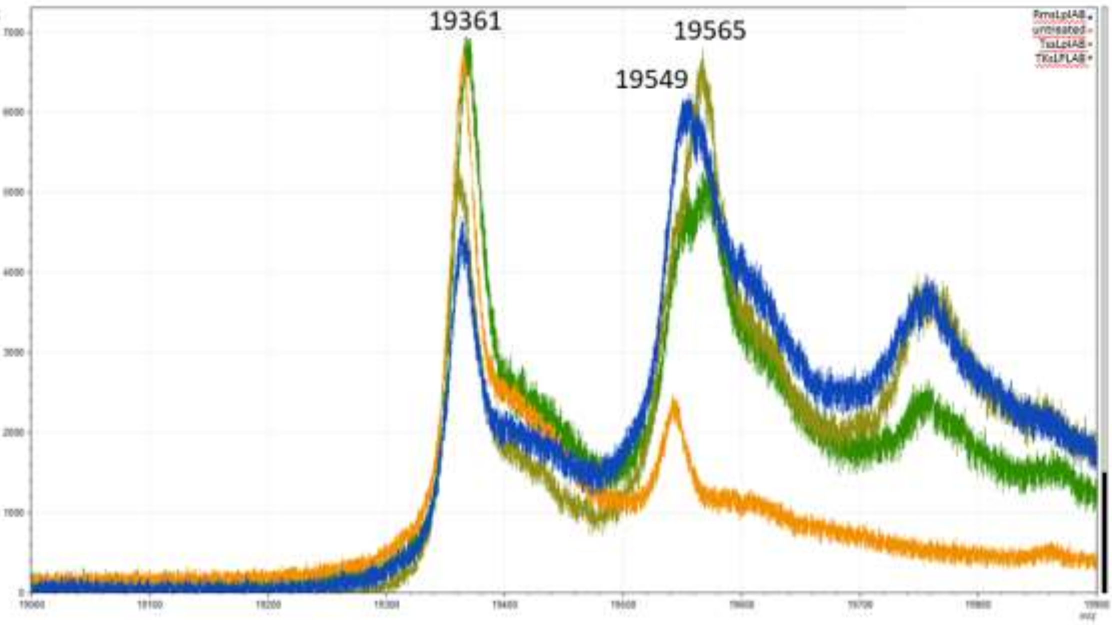


**Fig. S6. Mass spectrometry of *Aq*LbpA1_1 after incubation with different lipoate:protein ligases in the presence of 1.5 mM α-lipoic acid.** Recombinant *Aq*LbpA1_1 was already lipoylated to a very low extent. Lipoylation was catalyzed by all tested sLpl(AB)s. Note that the mass increase was 205 Da instead of the expected 189 Da in two reactions. This can be attributed to oxidation of the lipoyl group to its thiosulfinate form, resulting in modification of the target protein with a lipoyl residue carrying one additional oxygen atom (+16 Da) (Krishnan & Garnett, 2011; Reed, DeBusk, Gunsalus, & Hornberger Jr., 1951; Reed, Gunsalus et al., 1953; Stary, Jindal, & Murray, 1975).

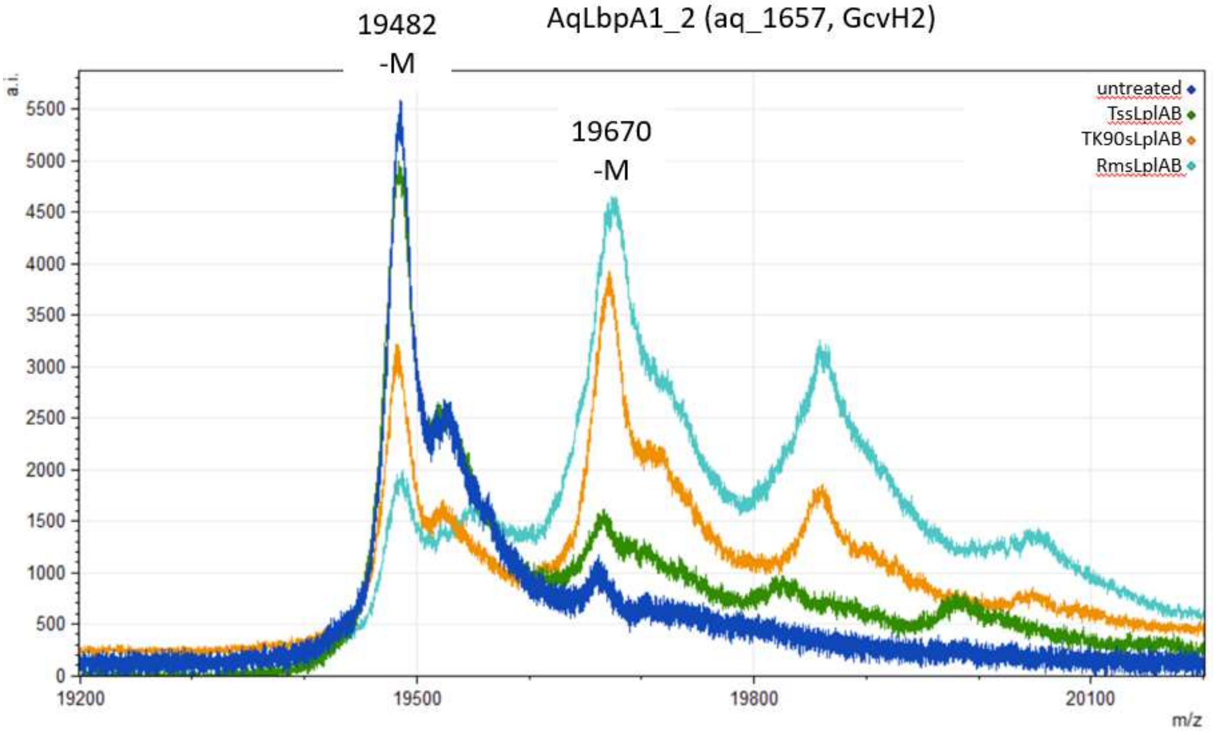


**Fig. S7. Mass spectrometry of *Aq*LbpA1_2 after incubation with different lipoate:protein ligases in the presence of 1.5 mM α-lipoic acid.** Lipoylation was catalyzed by *Rm*sLpl(AB) and *TK90*sLpl(AB), while *Tss*Lpl(AB) proved unable to modify the target. –M, the detected mass corresponds to the full-length polypeptide lacking the initiator methionine.

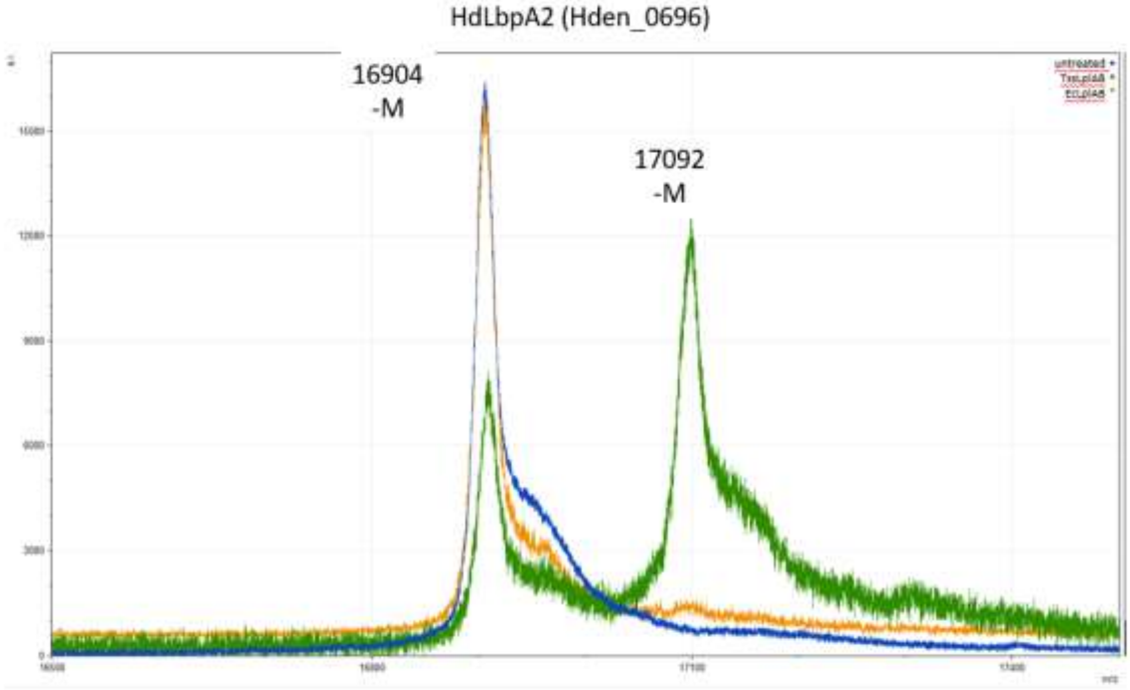


**Fig. S8. Mass spectrometry of *Hd*LbpA2 after incubation with different lipoate:protein ligases in the presence of 1.5 mM α-lipoic acid.** Efficient lipoylation was catalyzed by sLpl(AB) from *Thiorhodospira sibirica,* whereas no modification was observed in the presence of *Ec*Lpl(AB). –M, the detected mass corresponds to the full-length polypeptide lacking the initiator methionine.

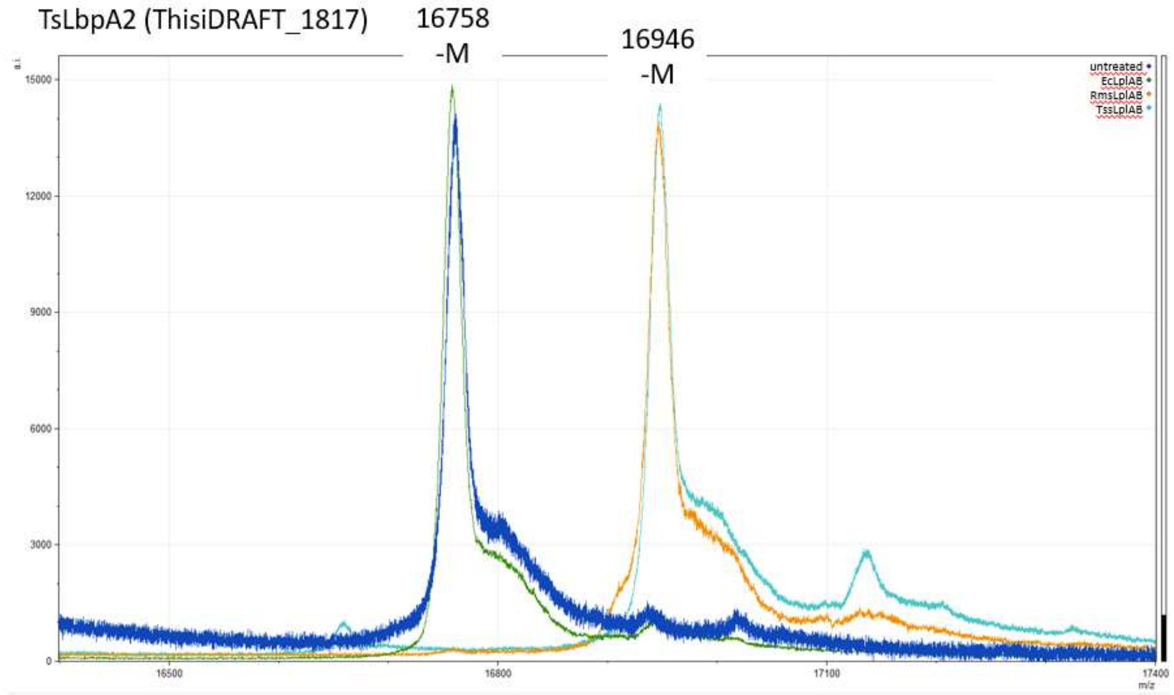


**Fig. S9. Mass spectrometry of *Ts*LbpA2 after incubation with different lipoate:protein ligases in the presence of 1.5 mM α-lipoic acid.** Efficient lipoylation was catalyzed by sLpl(AB) from *Thiorhodospira sibirica* and *Roseovarius mucosus*, whereas no modification was observed in the presence of *Ec*Lpl(AB). –M, the detected mass corresponds to the full-length polypeptide lacking the initiator methionine.

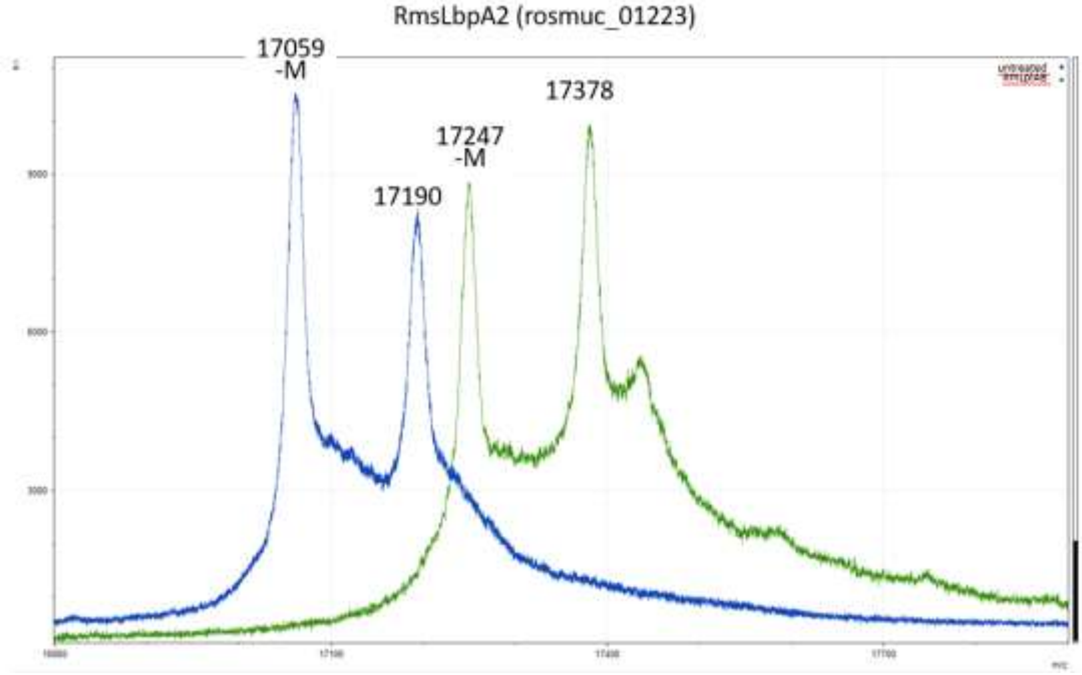


**Fig. S10. Mass spectrometry of RmLbpA2 after incubation with different lipoate:protein ligases in the presence of 1.5 mM α-lipoic acid.** Efficient lipoylation was catalyzed by sLpl(AB) from *Thiorhodospira sibirica* and *Roseovarius mucosus*, whereas no modification was observed in the presence of *Ec*Lpl(AB). –M, the detected mass corresponds to the full-length polypeptide lacking the initiator methionine.

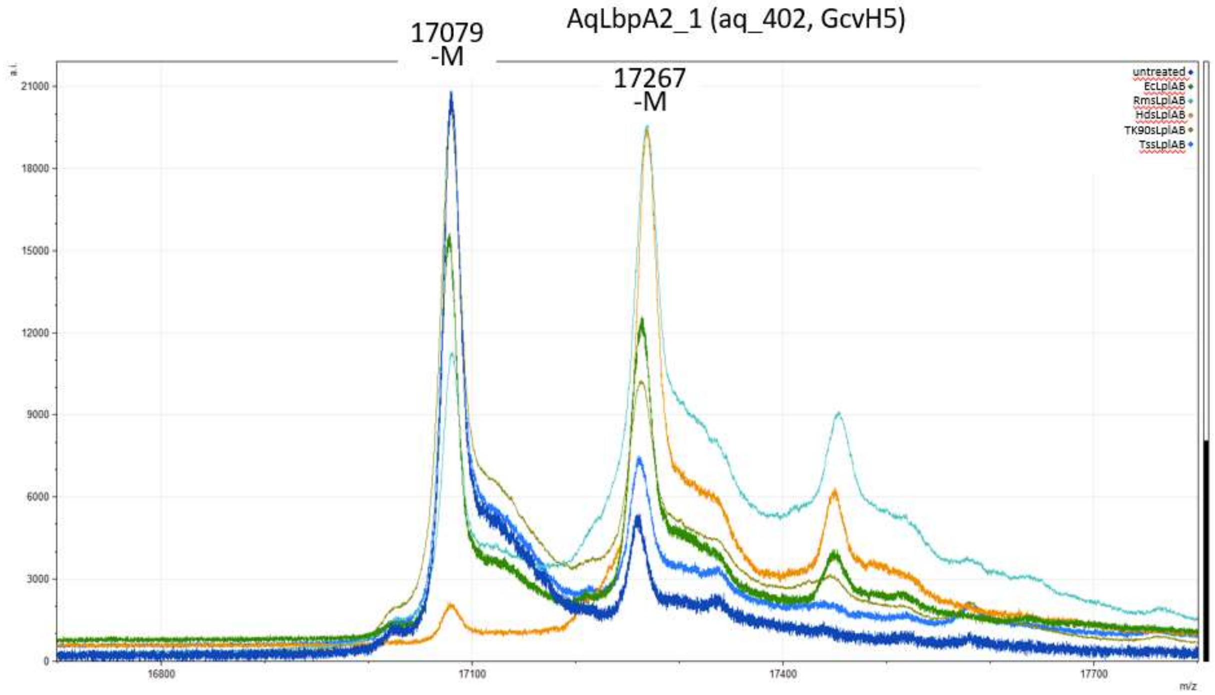


**Fig. S11. Mass spectrometry of *Aq*LbpA2_1 after incubation with different lipoate:protein ligases in the presence of 1.5 mM α-lipoic acid.** Recombinant AqLbpA2_1 was already lipoylated to a very low extent. Complete lipoylation was catalyzed by sLpl(AB) from *H. denitrificans*, whereas lipoylation by *Ec*Lpl(AB) and *Rms*Lpl(AB) was much less efficient. The degree of lipoylation hardly increased upon incubation with TK90sLpl(AB) or *Ts*sLpl(AB). –M, the detected mass corresponds to the full-length polypeptide lacking the initiator methionine.

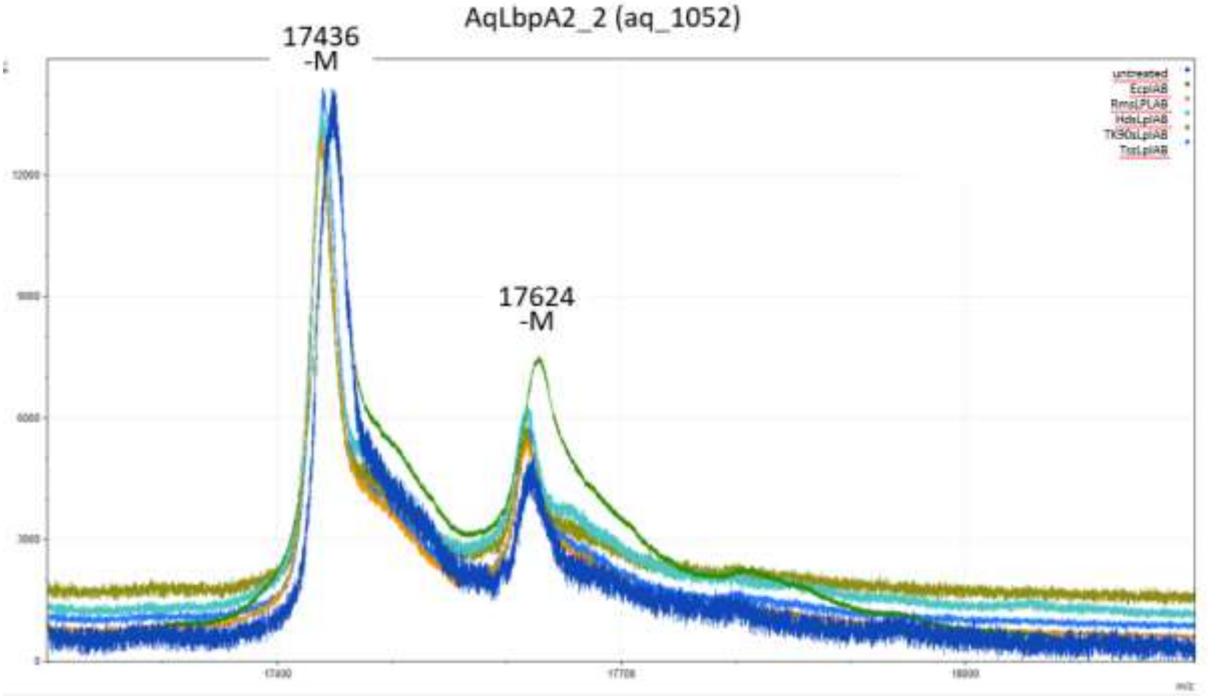


**Fig. S12. Mass spectrometry of *Aq*LbpA2_2 after incubation with different lipoate:protein ligases in the presence of 1.5 mM α-lipoic acid.** Recombinant *Aq*LbpA2_2 was already lipoylated to a small extent. Incubation with the various ligases did not result in any further lipoylation. –M, the detected mass corresponds to the full-length polypeptide lacking the initiator methionine.

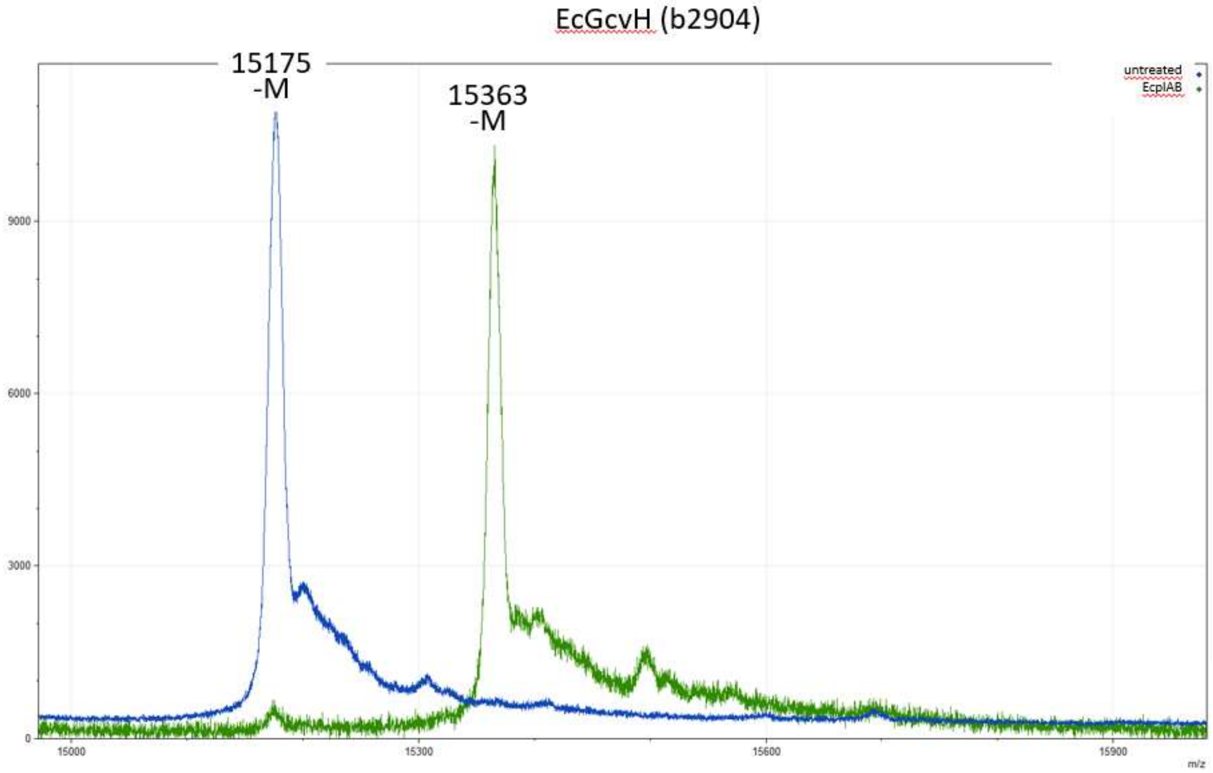


**Fig. S13. Mass spectrometry of *Ec*GcvH after incubation with lipoate:protein ligase from *E. coli* in the presence of 1.5 mM α-lipoic acid.**–M, the detected mass corresponds to the full-length polypeptide lacking the initiator methionine.

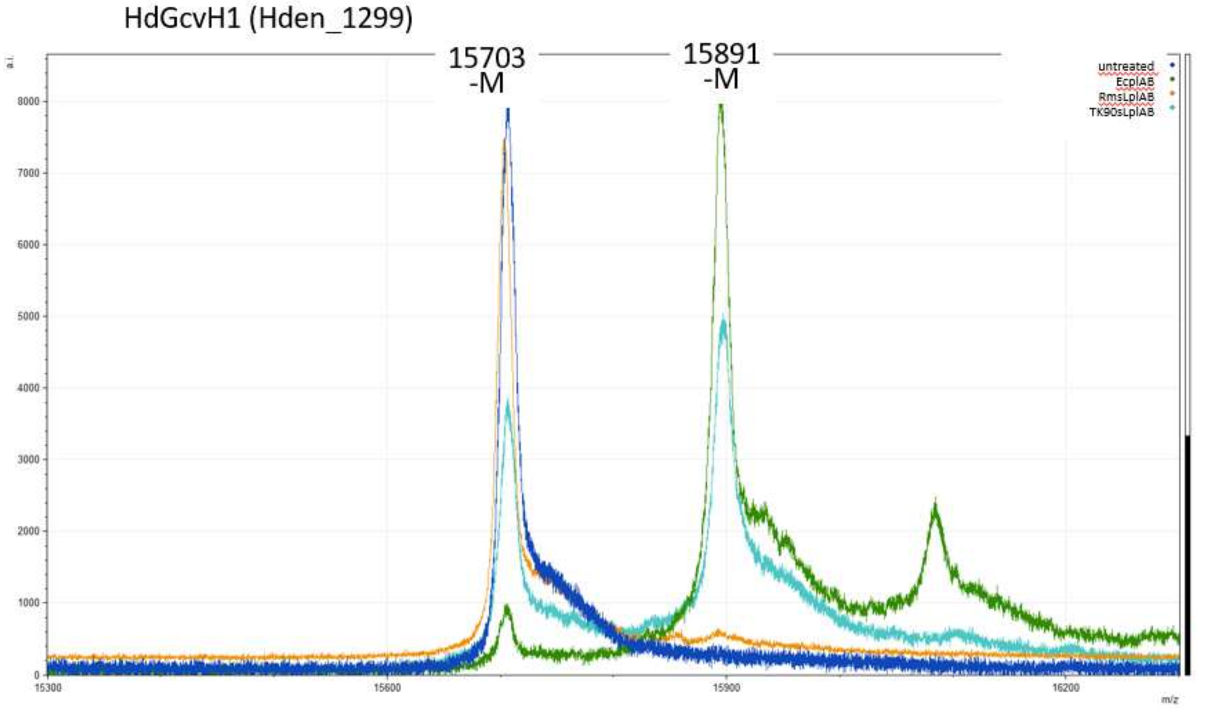


**Fig. S14. Mass spectrometry of *Hd*GcvH_1 after incubation with different lipoate:protein ligases in the presence of 1.5 mM α-lipoic acid.** *Hd*GcvH_1 was efficiently lipoylated by *Ec*Lpl(AB). *TK90*sLpl(AB) also catalyzed lipoylation of the target, albeit to a much lower extent. *Rm*sLpl(AB) did not accept *Hd*GcvH_1 as a substrate. –M, the detected mass corresponds to the full-length polypeptide lacking the initiator methionine.

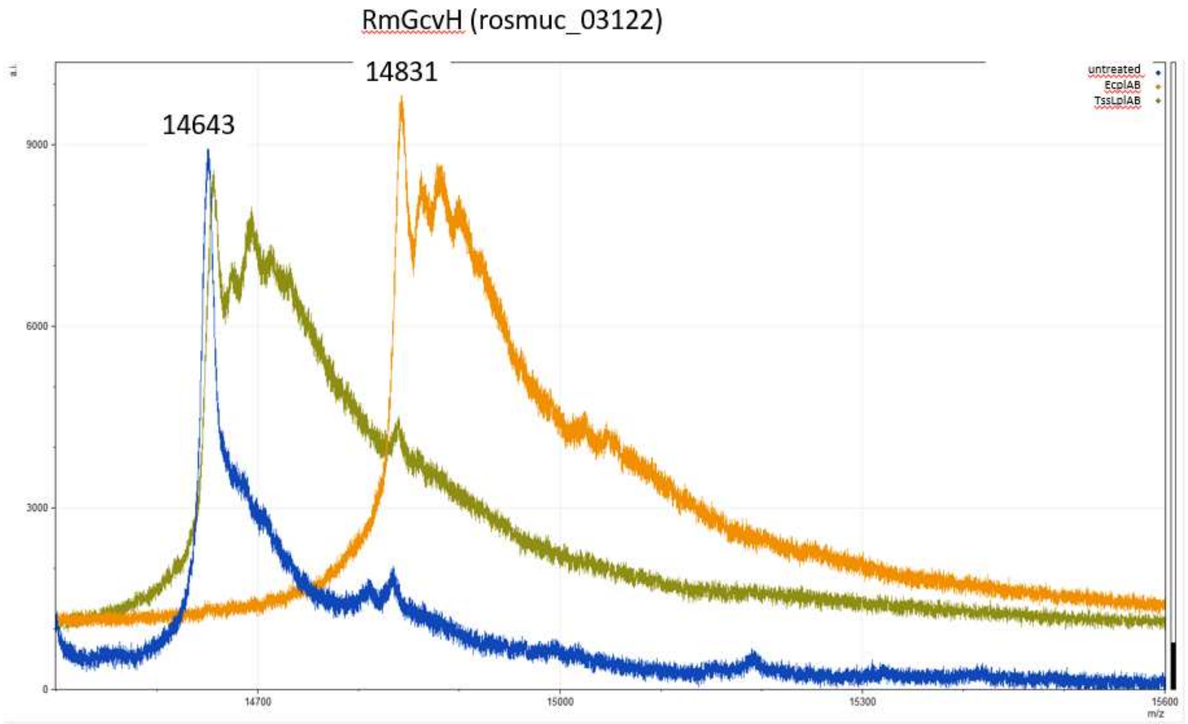


**Fig. S15. Mass spectrometry of *Rm*GcvH after incubation with different lipoate:protein ligases in the presence of 1.5 mM α-lipoic acid.** *Rm*GcvH_1 was efficiently lipoylated by *Ec*Lpl(AB). *Ts*sLpl(AB) did not accept *Rm*GcvH as a substrate. –M, the detected mass corresponds to the full-length polypeptide lacking the initiator methionine.

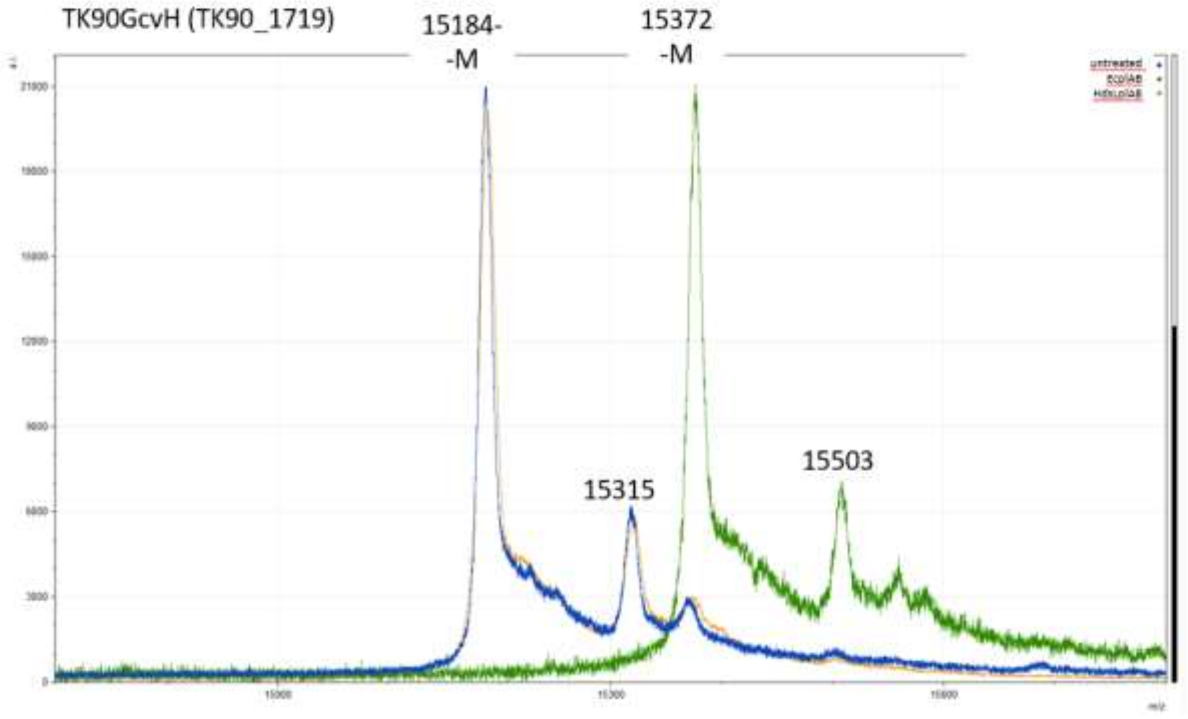


**Fig. S16. Mass spectrometry of *TK90*GcvH after incubation with different lipoate:protein ligases in the presence of 1.5 mM α-lipoic acid.** *TK90*GcvH was efficiently lipoylated by *Ec*Lpl(AB). *Hd*sLpl(AB) did not accept *TK90*GcvH as a substrate. –M, the detected mass corresponds to the full-length polypeptide lacking the initiator methionine.

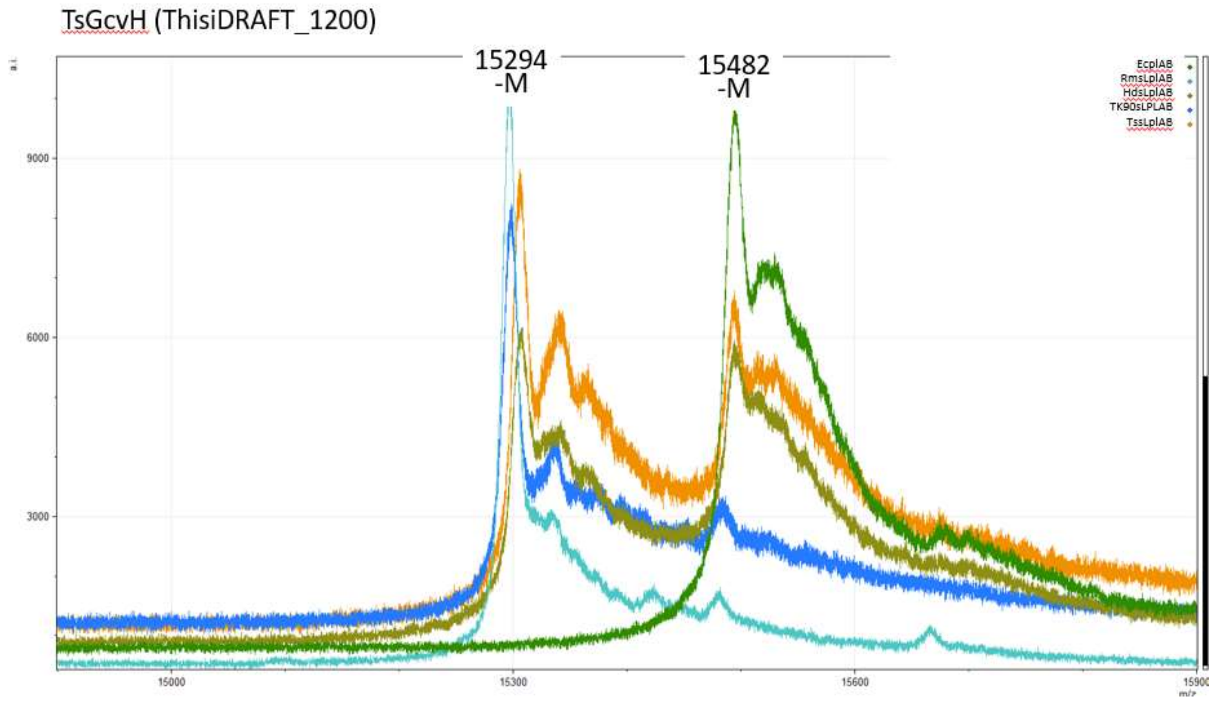


**Fig. S17. Mass spectrometry of *Ts*GcvH after incubation with different lipoate:protein ligases in the presence of 1.5 mM α-lipoic acid.** *Ts*GcvH was completely lipoylated by *Ec*Lpl(AB). *TK90*sLpl(AB), *Hd*sLpl(AB) and *Ts*sLpl(AB) also catalyzed lipoylation of the target, albeit to a very low extent. *Rm*sLpl(AB) did not accept *Ts*GcvH as a substrate. –M, the detected mass corresponds to the full-length polypeptide lacking the initiator methionine.

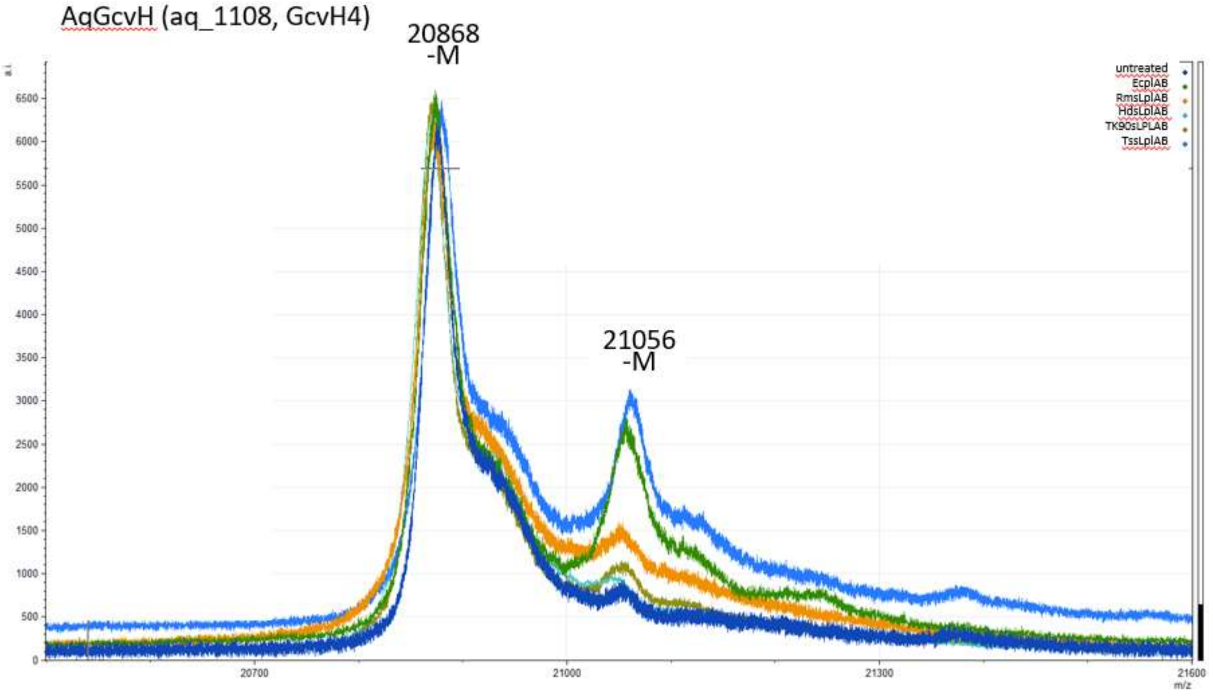


**Fig. S18. Mass spectrometry of *Aq*GcvH after incubation with different lipoate:protein ligases in the presence of 1.5 mM α-lipoic acid.** *Aq*GcvH was partially lipoylated by *Ec*Lpl(AB) and *Ts*sLpl(AB). The other sLpl(AB) enzymes did not accept *Aq*GcvH as a substrate. –M, the detected mass corresponds to the full-length polypeptide lacking the initiator methionine.

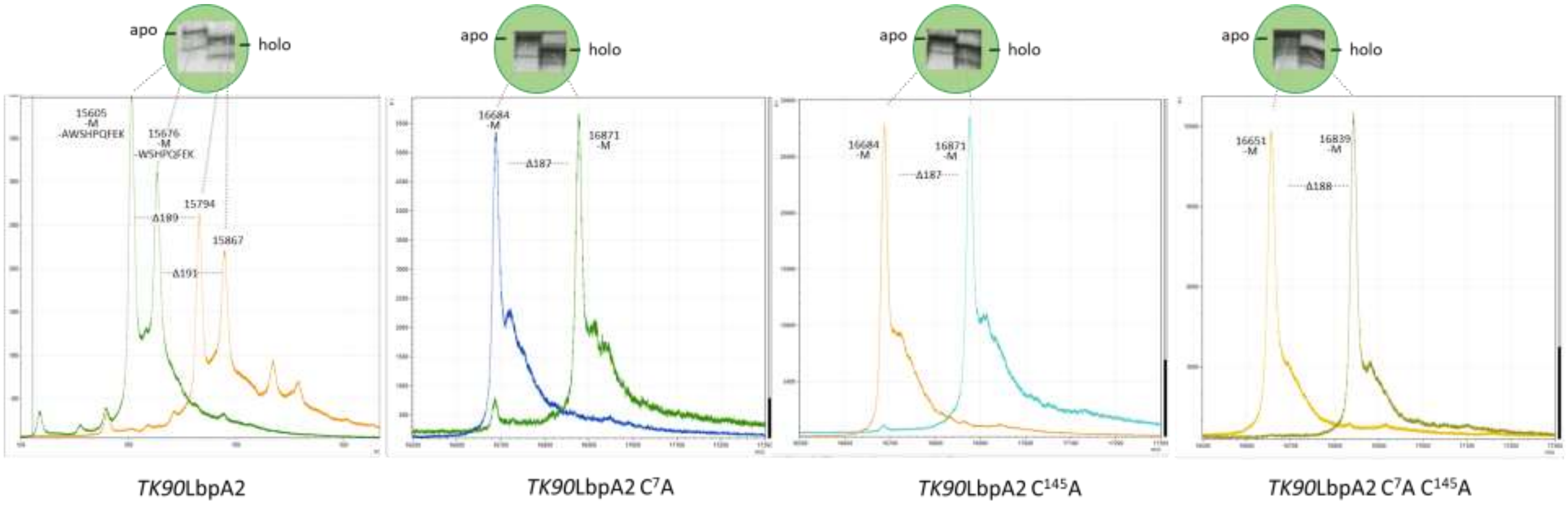


**Fig. S19. Gel shift assays and mass spectrometric analysis of *TK90*LbpA2 and Cys-to-Ala variants after incubation with *TK90*sLpl(AB) in the presence of 1.5 mM α-lipoic acid.** All four protein variants were efficiently lipoylated, as evidenced by the faster migration of the holo-proteins in native PAGE and mass increases consistent with the addition of a lipoyl moiety (+189 Da) –M, indicates that the detected mass corresponds to the full-length polypeptide lacking the initiator methionine. *TK90*LbpA2 was detected exclusively in two truncated forms, lacking either the C-terminal Strep-tag or the Strep-tag together with an additional alanine residue.

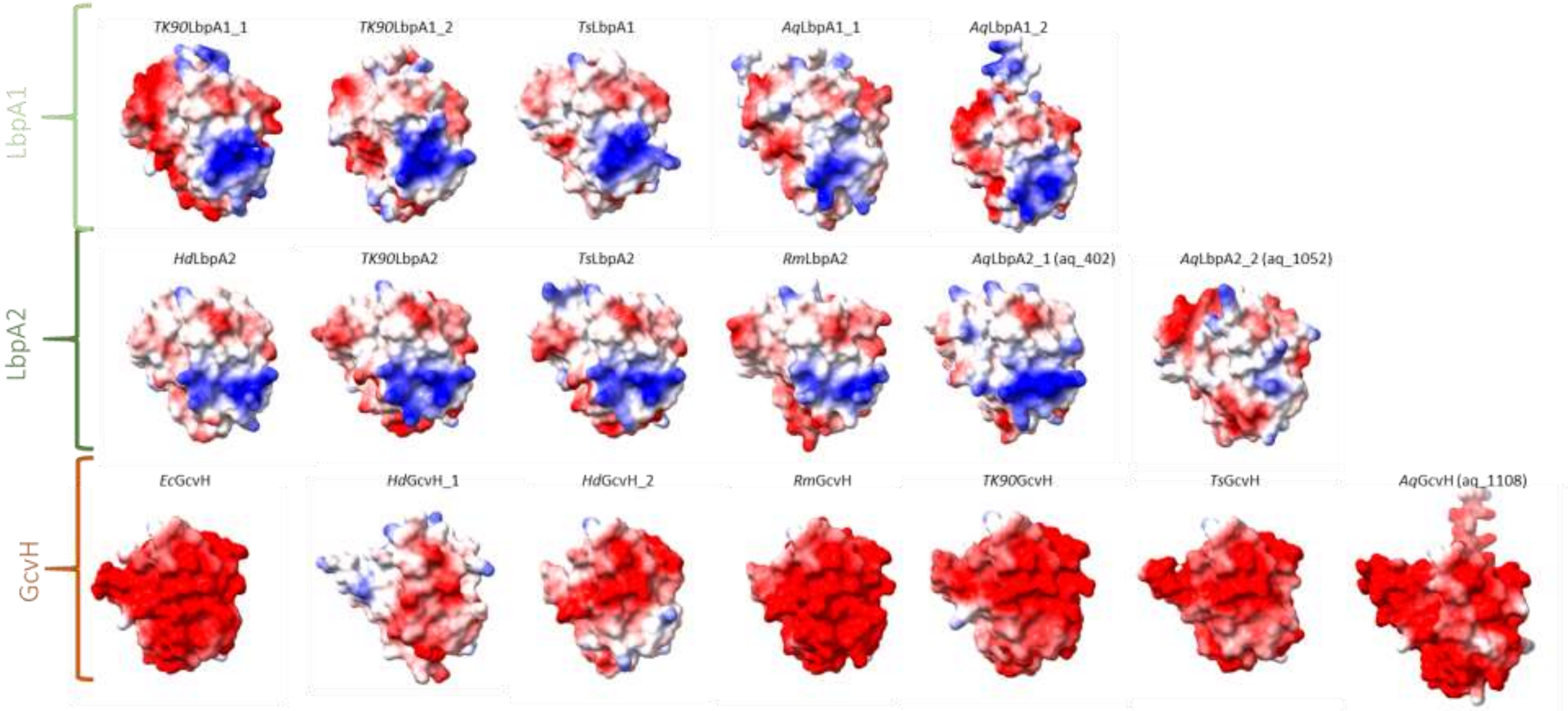


**Fig. S20. Structures of bacterial LbpA1, LbpA2 and GcvH proteins.** Except *Ec*GcvH, for which an experimentally determined crystal structure is available (PDB 3AB9 (Okamura-Ikeda, Hosaka et al., 2010), all structures were predicted by Alphafold (Jumper, Evans et al., 2021; Varadi, Bertoni et al., 2024). Structures are colored according to their surface electrostatic potential: red, acidic; blue, basic; white, neutral. All proteins are displayed in the same orientation.

**Table S2: Strains and plasmids**

| Strains or plasmids | Relevant genotype, description | Reference or source |
|---|---|---|
| **Strains** | | |
| *E. coli* 10-beta | Δ(*ara-leu*) 7697 *araD*139 *fhuA ΔlacX74 galK16 galE15 e14-* *ϕ80dlacZΔM15 recA1 relA1 endA1* *nupG rpsL* ($Str^R$) *rph spoT1* Δ(*mrr-hsdRMS-mcrBC*) | New England Biolabs |
| *E. coli* DH5α | $F^-$ φ80*lacZ*ΔM15 Δ(*lacZYA-argF*)U169 *recA1 end*A1 *hsdR17*($r_K^-$, $m_K^+$) phoA *supE44* λ–*thi*-1 *gyrA*96 *rel*A1 | New England Biolabs |
| *E. coli* BL21 (DE3) | $F^-$ *ompT hsdS*$_B$ ($r_B^-$, $m_B^-$) *gal dcm* (DE3) | Novagen |
| *E. coli* K12, MG1655, DSM 18039 | Wild type | DSMZ |
| *Hyphomicrobium denitrificans* DSM 1869$^T$ | Wild type | (Urakami, Sasaki, Suzuki, & Komagata, 1995) |
| *Thioalkalivibrio* sp. K90mix | Wild type | (Muyzer, Sorokin et al., 2011) |
| *Thiorhodospira sibirica*. ATCC 700588$^T$ | Wild type | (Bryantseva, Gorlenko et al., 1999) |
| *Roseovarius mucosus* DSM 17069$^T$ | Wild type | (Biebl, Allgaier et al., 2005; Riedel, Spring et al., 2015) |
| *Aquifex aeolicus* VF5$^T$ | Wild type | (Deckert, Warren et al., 1998) |
| **Plasmids** | | |
| pET22b(+) | $Amp^R$, T7 promoter, lac operator, C-terminal His tag, pelB leader | Novagen |
| pET28a(+) | $Km^R$, T7 promoter, lac operator, C-terminal His tag, pelB leader | Novagen |
| pET22b(+)_Strep-M | $Amp^R$, T7 promoter, lac operator, C-terminal Strep-tag, Strep-tag encoding sequence inserted as a NdeI-XhoI PCR fragment generated with primers Fwd_pET22b-Strep and Rev_pET22b-Strep using pET22b(+) as a template | This work |
| pET28a(+)_*Ec*Lpl(AB)_N-His | $Km^R$, N-terminal His-tag, NdeI-XhoI fragment of PCR amplified *lpl*A (b4386) in pET28a(+) | This work |
| pET22b(+)_*TK90*sLpl(AB)_C-Strep | $Amp^R$, C-terminal His-tag, NdeI-HindIII fragment of PCR amplified *slpl(AB)* (TK90_0642) in pET22b(+) | This work |
| pET28a(+)_*Ts*sLpl(AB)_C-His | $Km^R$, C-terminal His-tag, NcoI-HindIII fragment of PCR amplified *slpl(AB)* (ThisiDRAFT_2298) in pET28a(+) | This work |
| pET22b(+)_*Hd*sLpl(AB)_C-His | $Amp^R$, C-terminal His-tag, NdeI-XhoI fragment of PCR amplified *slpl(AB)* (Hden_0686) in pet22b(+) | This work |
| pET28a(+)_*Rm*sLpl(AB)_C-His | $Km^R$, C-terminal His-tag, NcoI-NotI fragment of PCR amplified *slpl(AB)* (rosmuc_01225) in pET28a(+) | This work |
| pET28a(+)_*TK90*sLpl(AB)_2_N-His | $Km^R$, N-terminal His-tag, NdeI-HindIII fragment of PCR amplified *slpl(AB)_2* (TK90_0648) in in pET28a(+) | This work |
| pET28a(+)_*Aq*LbpA1_1_N-Strep | $Km^R$, N-terminal Strep-tag, gene of AqLbpA1_1 (aq_944) codon-optimized for *E. coli* and cloned into NcoI/HindIII sites of pET28a(+), purchased from BioCat GmbH, Heidelberg, Germany | This work |
| pET28a(+)_*Aq*LbpA1_2_N-Strep | $Km^R$, N-terminal Strep-tag, gene of AqLbpA1_2 (aq_1657) codon-optimized for *E. coli* and cloned into NcoI/HindIII sites of pET28a(+), purchased from BioCat GmbH, Heidelberg, Germany | This work |
| pET28a(+)_*Aq*LbpA2_1_N-Strep | $Km^R$, N-terminal Strep-tag, gene of AqLbpA2_1 (aq_402) codon-optimized for *E. coli* and cloned into NcoI/HindIII sites of pET28a(+), purchased from BioCat GmbH, Heidelberg, Germany | This work |

| | | |
|---|---|---|
| pET28a(+)_*Aq*LbpA2_2_N-Strep | Km$^R$, N-terminal Strep-tag, gene of AqLbpA2_2 (aq_1052) codon-optimized for *E. coli* and cloned into NcoI/HindIII sites of pET28a(+), purchased from BioCat GmbH, Heidelberg, Germany | This work |
| pET28a(+)_*Aq*GcvH_N-Strep | Km$^R$, N-terminal Strep-tag, gene of AqGcvH (aq_1108) codon-optimized for *E. coli* and cloned into NcoI/HindIII sites of pET28a(+), purchased from BioCat GmbH, Heidelberg, Germany | This work |
| pET22b(+)_TK90_LbpA1_1_C-Strep | Amp$^R$, C-terminal Strep-tag, NdeI-HindIII fragment of PCR amplified *lbp*A1_1 (TK90_0638) in pET22b(+)_Strep-M | This work |
| pET22b(+)_*TK90*LbpA1_2_C-Strep | Amp$^R$, C-terminal Strep-tag, NdeI-HindIII fragment of PCR amplified *lbp*A1_2 (TK90_1746) in pET22b(+)_Strep-M | This work |
| pET22b(+)_*TK90*LbpA2_C-Strep | Amp$^R$, C-terminal Strep-tag, NdeI-HindIII fragment of PCR amplified *lbp*A2 (TK90_0640) in pET22b(+)_Strep-M | This work |
| pET22b(+)_*TK90*GcvH_C-Strep | Amp$^R$, C-terminal Strep-tag, NdeI-HindIII fragment of PCR amplified *gcvH* (TK90_1719) in pET22b(+)_Strep-M | This work |
| pET22b(+)_ThisiLbpA1_C-Strep | Amp$^R$, C-terminal Strep-tag, NdeI-HindIII fragment of PCR amplified *lbpA1* (ThisiDRAFT_0958) in pET22b(+)_Strep-M | This work |
| pET22b(+)_ThisiLbpA2_C-Strep | Amp$^R$, C-terminal Strep-tag, NdeI-HindIII fragment of PCR amplified *lbpA2* (ThisiDRAFT_1817) in pET22b(+)_Strep-M | This work |
| pET22b(+)_ThisiGcvH_C-Strep | Amp$^R$, C-terminal Strep-tag, NdeI-HindIII fragment of PCR amplified *gcvH* (ThisiDRAFT_1200) in pET22b(+)_Strep-M | This work |
| pET22b(+)_RmLbpA2_C-Strep | Amp$^R$, C-terminal Strep-tag, NdeI-HindIII fragment of PCR amplified *lbpA2* (rosmuc_01223) in pET22b(+)_Strep-M | This work |
| pET22b(+)_RmGcvH_C-Strep | Amp$^R$, C-terminal Strep-tag, NdeI-HindIII fragment of PCR amplified *gcvH* (rosmuc_03122) in pET22b(+)_Strep-M | This work |
| pET22b(+)_HdenLbpA2_C-Strep | Amp$^R$, C-terminal Strep-tag, NdeI-HindIII fragment of PCR amplified *lbpA2* (Hden_0696) in pET22b(+)_Strep-M | This work |
| pET22b(+)_Hden_GcvH1_C-Strep | Amp$^R$, C-terminal Strep-tag, NdeI-HindIII fragment of PCR amplified *gcvH1* (Hden_1299) in pET22b(+)_Strep-M | This work |
| pET22b(+)_Hden_GcvH2_C-Strep | Amp$^R$, C-terminal Strep-tag, NdeI-HindIII fragment of PCR amplified *gcvH2* (Hden_2793) in pET22b(+)_Strep-M | This work |
| pET22b(+)_EcGcvH_C-Strep | Amp$^R$, C-terminal Strep-tag, NdeI-HindIII fragment of PCR amplified *gcvH* (b2904) in pET22b(+)_Strep-M | This work |
| pET22b(+)_*TK90*LbpA2_C-Strep_C7A | Amp$^R$, C-terminal Strep-tag, pET22b(+)_*Tk90*LbpA2_C-Strep with Cys$^7$Ala exchange | This work |
| pET22b(+)_*TK90*LbpA2_C-Strep_C145A | Amp$^R$, C-terminal Strep-tag, pET22b(+)_*Tk90*LbpA2_C-Strep with Cys$^{145}$Ala exchange | This work |
| pET22b(+)_*TK90*LbpA2_C-Strep_C7A_C145A | Amp$^R$, C-terminal Strep-tag, pET22b(+)_*Tk90*LbpA2_C-Strep with Cys$^7$Ala and Cys$^{145}$Ala exchanges | This work |
| pET22b(+)_RmLbpA2_TPKR51-54DLPE-C-Strep | Amp$^R$, C-terminal Strep-tag, pET22b(+)_*Rm*LbpA2_C-Strep (rosmuc_01223) with TPKR$^{51-54}$DLPE exchange | This work |
| pET22b(+)_RmLbpA2_R63A-C-Strep | Amp$^R$, C-terminal Strep-tag, pET22b(+)_*Rm*LbpA2_C-Strep (rosmuc_01223) with Arg$^{63}$Ala exchange | This work |

| | | |
|---|---|---|
| pET22b(+)_RmLbpA2_ P76D-C-Strep | Amp[R], C-terminal Strep-tag, pET22b(+)_*Rm*LbpA2_C-Strep (rosmuc_01223) with Pro$^{76}$Asp exchange | This work |
| pET28a(+)_RmLbpA2_ mutagenesis12 | Km[R], C-terminal Strep-tag, pET28a(+)_*Rm*LbpA2_C-Strep (rosmuc_01223) with TPKR$^{51-54}$DLPE and R$^{63}$A exchanges, codon-optimized for *E. coli* and cloned into NdeI/HindIII sites of pET28a(+), purchased from BioCat GmbH, Heidelberg, Germany | This work |
| pET28a(+)_RmLbpA2_mutagenesis13 | Km[R], C-terminal Strep-tag, pET28a(+)_*Rm*LbpA2_C-Strep (rosmuc_01223) with TPKR$^{51-54}$DLPE and P$^{76}$D exchanges, codon-optimized for *E. coli* and cloned into NdeI/HindIII sites of pET28a(+), purchased from BioCat GmbH, Heidelberg, Germany | This work |
| pET28a(+)_RmLbpA2_mutagenesis23 | Km[R], C-terminal Strep-tag, pET28a(+)_*Rm*LbpA2_C-Strep (rosmuc_01223) with R$^{63}$A and P$^{76}$D exchanges, codon-optimized for *E. coli* and cloned into NdeI/HindIII sites of pET28a(+), purchased from BioCat GmbH, Heidelberg, Germany | This work |
| pET28a(+)_RmLbpA2_ mutagenesis123 | Km[R], C-terminal Strep-tag, pET28a(+)_*Rm*LbpA2_C-Strep (rosmuc_01223) with TPKR$^{51-54}$DLPE, R$^{63}$A and P$^{76}$D exchanges, codon-optimized for *E. coli* and cloned into NdeI/HindIII sites of pET28a(+), purchased from BioCat GmbH, Heidelberg, Germany | This work |

**Table S2: Primers**

| Primer | Sequence | Reference or source |
|---|---|---|
| rosmuc01223_NdeI_fw | TAT**CATATG**GTTTCTGTCAAGGGCTGCG (NdeI) | This work |
| rosmuc01223_HindIII_rev | ATA**AAGCTT**CTGACAGCCCGGAAAATCAT (HindIII) | This work |
| rosmuc01225_NcoI_fw | TAT**CCATGG** GT ACACCCTTTCGCCTGATAG (NcoI) | This work |
| rosmuc01225_NotI_rev | ATA**GCGGCCGC**TGATCTGGCATCCTCAGAAA (NotI) | This work |
| rosmuc03122_NdeI_fw | TAT**CATATG**AAATTCACCGAAGAACATGAATGG (NdeI) | This work |
| rosmuc03122_HindIII_rev | ATA**AAGCTT**GGCGATATGGGCCTGATA (HindIII) | This work |
| TK90-0638-NdeI-fw | GCG**CATATG**GACTGCAACGGTTGCGAGTTTC (NdeI) | This work |
| TK90-0638-HindIII-rev | CCGG**AAGCTT**TCGCAGCTTGCGCTGACACTCGATG (HindIII) | This work |
| TK90-0640-NdeI-fw | CTT**CATATG**GGCGCAGTACGGGGTTGTGATATTC (NdeI) | This work |
| TK90-0640-HindIII-rev | CCGG**AAGCTT**TCGCAGCTTGCGCTGACACTCGATG (HindIII) | This work |
| TK90_0642_fw | TAT**CATATG**TCTGAACGCGATCAGAAGACG (NdeI) | This work |
| TK90_0642_rev | ATA**AAGCTT**TCCCGCTGATTCCTCGGT (HindIII) | This work |
| TK90_0648_NdeI_fw | TAT**CATATG**TTGGTCGCATCCACGGTCACC (NdeI) | This work |
| TK90_0648_HindIII_rev | ATA**AAGCTT** TCA GGCGGCGGCGATC (HindIII) | This work |
| TK90_1719_NdeI_fw | TAT**CATATG**AGCGAAGCACCGAAGGATCT (NdeI) | This work |
| TK90_1719_HindIII_rev | ATA**AAGCTT**GGACTCGGCCTCGACGT (HindIII) | This work |
| TK90_1746_NdeI_fw | TAT**CATATG**AGCCATACCGAATACAACGGCT (NdeI) | This work |
| TK90_1746_HindIII_rev | ATA**AAGCTT**GTCGGCACAGCGCATG (HindIII) | This work |
| ThisiDRAFT1817_Nde_fw | CTGG**CATATG**GGCACTGCAAAAGGCTGTC (NdeI) | This work |
| ThisiDRAFT1817_HindIII_rev | ATAC**AAGCTT**AGCCCCGCACCCCTTAAAG (HindIII) | This work |
| ThisiDRAFT0958_Nde_fw | GCCG**CATATG**GATGACTGTAACGGCTGTGAATTTC (NdeI) | This work |
| ThisiDRAFT0958_HindIII_rev | AATG**AAGCTT**TGCCGTGCGGGTGCAGTGG (HindIII) | This work |
| ThisiDRAFT1200_NdeI_fw | TAT**CATATG**AGCGATTTTCCTGCTGAATTACGC (NdeI) | This work |
| ThisiDRAFT1200_HindIII_rev | ATA**AAGCTT**GCTATCATCGGCGTCAAGGC (HindIII) | This work |
| ThisiDRAFT2298_NcoI_Fw | TAT**CCATGG**GTACCCGACATTCCCAAACGCT (NcoI) | This work |
| ThisiDRAFT2298_HindIII_rev | ATA**AAGCTT**AGATGATGCCGCCACTTCTG (HindIII) | This work |
| Hden0686_NdeI_fw | GCGG**CATATG**GTGAGTATAGGGCGGCGC (NdeI) | This work |
| Hden0686_XhoI_rw | ATA**TCTCGAG**TGGCCTCAGCGTCCGCAAAG (XhoI) | This work |
| Hden0696_NdeI_fw | GTGGC**CATATG**CCAGTCGTGAAGGGATGC (NdeI) | This work |
| Hden0696_HindIII_rev | ACT**AAGCTT**CCAGCCGCAACCTGCGAAACC (HindIII) | This work |
| Hden1299_NdeI_fw | TAT**CATATG**CCCTCCAGCTTATCTGACGAGA (NdeI) | This work |
| Hden1299_HindIII_rev | ATA**AAGCTT**CCTGGCTTTTGCGCCGAG (HindIII) | This work |
| Hden2793_NdeI_fw | TAT**CATATG**CTGAAATATACCGAAGAGCATG (NdeI) | This work |
| Hden2793_NotI_rev | ATA**GCGGCCGC**AGCAATGAGGTTTGCGTAG (NotI) | This work |
| B2904_NdeI_fw | TAT**CATATG**AGCAACGTACCAGCAGAACT (NdeI) | This work |
| B2904_HindIII_rev | ATA**AAGCTT**CTCGTCTTCTAACAATGCTTCGT (HindIII) | This work |
| B4386_NcoI_fw | TAT**CCATGG** GT TCCACATTACGCCTGCTCATCT (NcoI) | This work |
| B4386_HindIII_rev | ATA**AAGCTT**CCTTACAGCCCCCGCC (HindIII) | This work |
| TK90_0640_C7A_fw | AGTACGGGGTGCGGATATTCCGGAAGATCTG | This work |

| | | |
|---|---|---|
| TK90_0640_C7A_rev | GCGCCCATATGTATATCTCCTT | This work |
| TK90_0640_C145A_fw | TTTCGGCGGCGCGAAGCTTGCGGCCGCG | This work |
| TK90_0640_C145A_rev | CCGTCCGCTTCCATCTTGGC | This work |
| RmLbpA2_TPKR$^{51-54}$DLPE_fw | CTTTGCTGTCGATCTGCCGGAAGTGGGGCGCCTGTTC | This work |
| RmLbpA2_TPKR$^{51-54}$DLPE_rev | ATCTGGCGGGATGCCAGTGC | This work |
| RmLbpA2_R63A_fw | AGGCAGGAGCGTCGGCGGCAACGATAG | This work |
| RmLbpA2_R63A_rev | CGAACAGGCGCCCCAC | This work |
| RmLbpA2_P76D_fw | ATGGGTGGGAGA ATGAGGCTTGCTTTCG | This work |
| RmLbpA2_P76D_rev | TTGCTACTTTCTATCGTTGCCGCC | This work |
| Fwd_pET22b-Strep | GAAGGAGATAT**ACATATG**AAAT (NdeI) | This work |
| Rev_pET22b-Strep | TTT**CTCGAG**TTATTTTTCGAACTGCGGGTGGCTCCACGCGGC CGCAAGCTTGTCGAC (XhoI) | This work |

### REFERENCES


Biebl, H., Allgaier, M., Lunsdorf, H., Pukall, R., Tindall, B. J., & Wagner-Dobler, I. (2005). *Roseovarius mucosus* sp. nov., a member of the Roseobacter clade with trace amounts of bacteriochlorophyll a. *International Journal of Systematic and Evolutionary Microbiology, 55*(Pt 6), 2377-2383.

Bryantseva, I. A., Gorlenko, V. M., Kompantseva, E. I., Imhoff, J. F., S�ling, J., & Mityushina, L. (1999). *Thiorhodospira sibirica* gen. nov., sp. nov., a new alkaliphilic purple sulfur bacterium from a Siberian soda lake. *International Journal of Systematic Bacteriology, 49*, 697-703.

Deckert, G., Warren, P. V., Gaasterland, T., Young, W. G., Lenox, A. L., Graham, D. E., Overbeek, R., Snead, M. A., Keller, M., Aujay, M., Huber, R., Feldman, R. A., Short, J. M., Olsen, G. J., & Swanson, R. V. (1998). The complete genome of the hyperthermophilic bacterium *Aquifex aeolicus*. *Nature, 392*(6674), 353-358.

Jumper, J., Evans, R., Pritzel, A., Green, T., Figurnov, M., Ronneberger, O., Tunyasuvunakool, K., Bates, R., Zidek, A., Potapenko, A., Bridgland, A., Meyer, C., Kohl, S. A. A., Ballard, A. J., Cowie, A., Romera-Paredes, B., Nikolov, S., Jain, R., Adler, J., Back, T., Petersen, S., Reiman, D., Clancy, E., Zielinski, M., Steinegger, M., Pacholska, M., Berghammer, T., Bodenstein, S., Silver, D., Vinyals, O., Senior, A. W., Kavukcuoglu, K., Kohli, P., & Hassabis, D. (2021). Highly accurate protein structure prediction with AlphaFold. *Nature, 596*(7873), 583-589.

Krishnan, C. V., & Garnett, M. (2011). Electrochemical behavior of the super antioxidant, α-lipoic acid. *International Journal of Electrochemical Science, 6*(8), 3607-3630.

Muyzer, G., Sorokin, D. Y., Mavromatis, K., Lapidus, A., Foster, B., Sun, H., Ivanova, N., Pati, A., D'Haeseleer, P., Woyke, T., & Kyrpides, N. C. (2011). Complete genome sequence of *Thioalkalivibrio* sp. K90mix. *Standards in Genomic Sciences, 5*(3), doi:10.4056/sigs.2315092.

Okamura-Ikeda, K., Hosaka, H., Maita, N., Fujiwara, K., Yoshizawa, A. C., Nakagawa, A., & Taniguchi, H. (2010). Crystal structure of aminomethyltransferase in complex with dihydrolipoyl-H-protein of the glycine cleavage system: implications for recognition of lipoyl protein substrate, disease-related mutations, and reaction mechanism. *Journal of Biological Chemistry, 285*(24), 18684-18692.

Reed, L. J., DeBusk, B. G., Gunsalus, I. C., & Hornberger Jr., C. S. (1951). Crystalline α-lipoic acid: a catalytic agent associated with pyruvate dehydrogenase. *Science, 114*(2952), 93-94.

Reed, L. J., Gunsalus, I. C., Schnakenberg, G. H. F., Soper, Q. F., Boaz, H. E., Kern, S. F., & Parke, T. V. (1953). Isolation, characterization and structure of α-lipoic acid. *Journal of the American Chemical Society, 75*(6), 1267-1270.

Riedel, T., Spring, S., Fiebig, A., Scheuner, C., Petersen, J., Goker, M., & Klenk, H. P. (2015). Genome sequence of the *Roseovarius mucosus* type strain (DSM 17069$^T$), a bacteriochlorophyll a-containing representative of the marine *Roseobacter* group isolated from the dinoflagellate *Alexandrium ostenfeldii*. *Standards in Genomic Sciences, 10*, 17.

Stary, F. E., Jindal, S. J., & Murray, R. W. (1975). Oxidation of $\alpha$-lipoic acid. *Journal of Organic Chemistry, 40*(1), 58-62.

Urakami, T., Sasaki, J., Suzuki, K. I., & Komagata, K. (1995). Characterization and description of *Hyphomicrobium denitrificans* sp. nov. *International Journal of Systematic Bacteriology, 45*(3), 528-532.

Varadi, M., Bertoni, D., Magana, P., Paramval, U., Pidruchna, I., Radhakrishnan, M., Tsenkov, M., Nair, S., Mirdita, M., Yeo, J., Kovalevskiy, O., Tunyasuvunakool, K., Laydon, A., Zidek, A., Tomlinson, H., Hariharan, D., Abrahamson, J., Green, T., Jumper, J., Birney, E., Steinegger, M., Hassabis, D., & Velankar, S. (2024). AlphaFold Protein Structure Database in 2024: providing structure coverage for over 214 million protein sequences. *Nucleic Acids Research, 52*(D1), D368-D375.
